# Activity-dependent refinement of nasal retinal projections drives topographic map sharpening in the teleost visual system

**DOI:** 10.1101/2020.12.14.422653

**Authors:** Olivia Spead, Fabienne E. Poulain

## Abstract

Topographic maps in the brain are essential for processing information. Yet, our understanding of topographic mapping has remained limited by our inability to observe maps forming and refining directly in vivo. Here, we used Cre-mediated recombination of a new colorswitch reporter in zebrafish to generate the first transgenic model allowing the dynamic analysis of retinotopic mapping in vivo. We found that the antero-posterior retinotopic map forms early but remains dynamic, with nasal and temporal retinal axons expanding their projection domains over time. Nasal projections initially arborize in the anterior tectum but progressively refine their projection domain to the posterior tectum in an activity-dependent manner. This activity-dependent refinement drives retinotopic map sharpening along the antero-posterior axis. Altogether, our study provides the first analysis of a topographic map maturing in real-time in a live animal and opens new strategies for dissecting the intricate mechanisms of precise topographic mapping in vertebrates.

## INTRODUCTION

Organization of neuronal connections into topographic maps is essential for the efficient transfer of information between brain regions. In the visual system, retinal projections transmit a precise and continuous representation of the external world by maintaining the neighboring relationship of the neurons they originate from in the retina (Cang and Feldheim, 2013; Huberman et al., 2008; McLaughlin and O’Leary, 2005; Triplett, 2014). Along the antero-posterior axis, retinal ganglion cells (RGCs) in the nasal retina project axons to the posterior tectum (or superior colliculus -SC- in mammals), whereas temporal RGCs innervate the anterior tectum. As Sperry first postulated in his chemoaffinity hypothesis (Sperry, 1963), studies in mouse, chick, xenopus and fish have demonstrated that this precise retinotopic map is first established by specific axon-target interactions, whereby axons with a specific set of receptors interpret guidance cues distributed in a gradient at the target. The repellent cues ephrin-As, for instance, are the main factors mediating topographic mapping along the antero-posterior axis. Subsequently to this guidance process, retinotopic projections are refined by activity-dependent mechanisms triggered by spontaneous waves of retinal activity or visual experience (Assali et al., 2014; Kutzarova et al., 2017; Leighton and Lohmann, 2016; Thompson et al., 2017).

It is now well accepted that both guidance cues at the target and patterned retinal activity act together in an instructive manner to establish a precise retinotopic map (Benjumeda et al., 2013; Cang et al., 2008; Pfeiffenberger et al., 2006). However, increasing evidence also implicates additional mechanisms in the establishment of retinotopy. Repulsive, competitive and stabilizing interactions among axons themselves are thought to play an important role in both initial mapping and refinement (Gosse et al., 2008; Hua et al., 2005; Louail et al., 2020; Rahman et al., 2020; Suetterlin et al., 2012; Suetterlin and Drescher, 2014; Spead and Poulain, 2020; Weth et al., 2014). Yet, our understanding of how and when trans-axonal signaling contributes to retinotopic map formation and maturation has remained limited by our inability to selectively manipulate RGCs in a topographic and reproducible manner. It also remains unclear how the retinotopic map becomes sharper as a whole, as we currently lack the ability to observe the map forming and refining over time in the same embryo in vivo. Fourier optical imaging of intrinsic signals allows for the visualization of functional collicular maps in the mouse but can only be employed once animals have developed vision, well after the retinotopic map has formed and refined (Cang et al., 2008). On the other hand, retinotopy can be analyzed at earlier stages by injecting lipophilic dyes or electroporating DNA plasmids in specific retinal quadrants. However, both approaches often require fixing specimen for analysis and have some degree of variability, which precludes the study of mapping dynamics and the detection of subtle topographic changes between times or conditions.

Because of its rapid external development and transparency, the zebrafish larvae has become a model of choice for the study of retinotopy (Förster et al., 2020; Kita et al., 2015; Poulain et al., 2010). Injections of lipophilic dyes in opposite regions of the retina have been used extensively to label retinotopic projections at different stages of development or regeneration and identify mutants with retinotopic defects (Baier et al., 1996; Harvey et al., 2019; Stuermer, 1988; Trowe et al., 1996; Xiao et al., 2005). Using that approach, early studies have shown that nasal and temporal retinal axons are localized at retinotopic sites as early as 3 days-post-fertilization (dpf), with temporal and nasal axons innervating the anterior and posterior tectum, respectively (Stuermer, 1988; Stuermer et al., 1990). Labeling a subset of RGCs in larvae fixed at 4 and 6 dpf has also revealed that projections cover a smaller territory at later stages, suggesting a refinement of the retinotopic map over time (Gnuegge et al., 2001). That reduced coverage is not observed in embryos treated with the voltage-gated sodium channel blocker tetrodotoxin (TTX), indicating a role for neuronal activity in retinotopic map maturation. While these observations highlight similar mechanisms underlying precise retinotopy in zebrafish and other species, we still do not know exactly when and how the retinotopic map refines and matures in teleosts. Advances in molecular genetics have allowed the generation of multiple transgenic lines for analyzing the lineage and functions of neuronal populations in zebrafish (Kawakami et al., 2016; Portugues et al.,2013; Robles, 2017), but the lack of enhancer driving transgene expression in specific retinal quadrants has precluded a similar unbiased analysis of retinotopic mapping over time in vivo.

Here, we report the generation of the first genetic model allowing the dynamic and quantitative analysis of retinotopic map formation and refinement directly in vivo. We show that an enhancer located upstream of *hmx1* and *hmx4* genes on chromosome 14 drives selective transgene expression in the nasal retina. We used Cre-mediated recombination of an *RGC:colorswitch* reporter to specifically label nasal and temporal retinal axons in vivo and image their projection domains at the tectum from 3 to 6 dpf by live confocal microscopy. Our analysis reveals that while the antero-posterior retinotopic map is formed at early developmental stages, it remains dynamic, with nasal and temporal axons expanding their projection domains over time. We further show that nasal retinal projections initially arborize in the anterior half of the tectum but progressively refine and condense their projection domain to the posterior tectum in an activity-dependent manner from 4 to 5 dpf. We finally demonstrate that the refinement of nasal projections drives the sharpening of the antero-posterior retinotopic map, and that both are prevented by genetically blocking neuronal activity in RGCs.

## RESULTS

### *hmx1* is expressed in the nasal retina throughout development

With the aim of identifying potential enhancers that would drive specific expression in the nasal or temporal retina throughout development in zebrafish, we first assessed genes that were previously described as selectively expressed in the nasal or temporal retinal half in vertebrates. Among them, the transcription factor *hmx1* has been specifically detected in the nasal retina of zebrafish, chick, and mice and was reported to regulate the retinal expression of EphA receptors (Boisset and Schorderet, 2012; Marcelli et al., 2014; Schulte and Cepko, 2000; Takahashi et al., 2009; Takahashi et al., 2003; Yoshiura et al., 1998). Given its role in retinal patterning, we decided to further analyze and quantify *hmx1* expression throughout retinotectal development by in situ hybridization (ISH). At 24 hpf, when optic cup morphogenesis is complete (Kwan et al., 2012), *hmx1* was strongly expressed in the nasal retina and the lens and was also faintly detected in the otic vesicle (**Figure 1A, A’**). At 48 hpf, when first retinal axons elongate along the tract and reach the tectum (Burrill and Easter, 1995; Stuermer, 1988), *hmx1* remained strongly expressed in the nasal retina and otic vesicle and could also be detected in the pharyngeal arches (**Figure 1B, B’, E**). Interestingly, while *hmx1* expression remained stable in the otic vesicle and pharyngeal arches over time, it became restricted to the RGC layer in the nasal retina at 72 hpf (**Figure 1C, C’, F**) and 96 hpf (**Figure 1D, D’, G**). Hmx1 expression could also be detected at lower levels in the nasal inner nuclear layer at both stages (**Figure 1F, G**). We further quantified the expression levels of *hmx1* along a 360° clockwise trajectory in the RGC layer at 48, 72, and 96 hpf (**Figure 1H**) and found a sharp gradient of expression along the nasal-temporal axis, with *hmx1* being highly expressed in the nasal half of the retina but absent from the temporal half at all three stages (**Figure 1I**). Since *hmx1* and its paralog *hmx4* arose from tandem duplication and are tightly linked on the same chromosome in chick and zebrafish (Adamska et al., 2001; Wotton et al., 2010), we also analyzed the expression of *hmx4* during retinal development (**Figure 1 - figure supplement 1**). As previously described in chick (Deitcher et al., 1994; Schulte and Cepko, 2000), *hmx4* had a similar expression to that of *hmx1* and was strongly detected in the nasal retina, otic vesicle and pharyngeal arches from 24 to 96 hpf.

**Figure 1.**
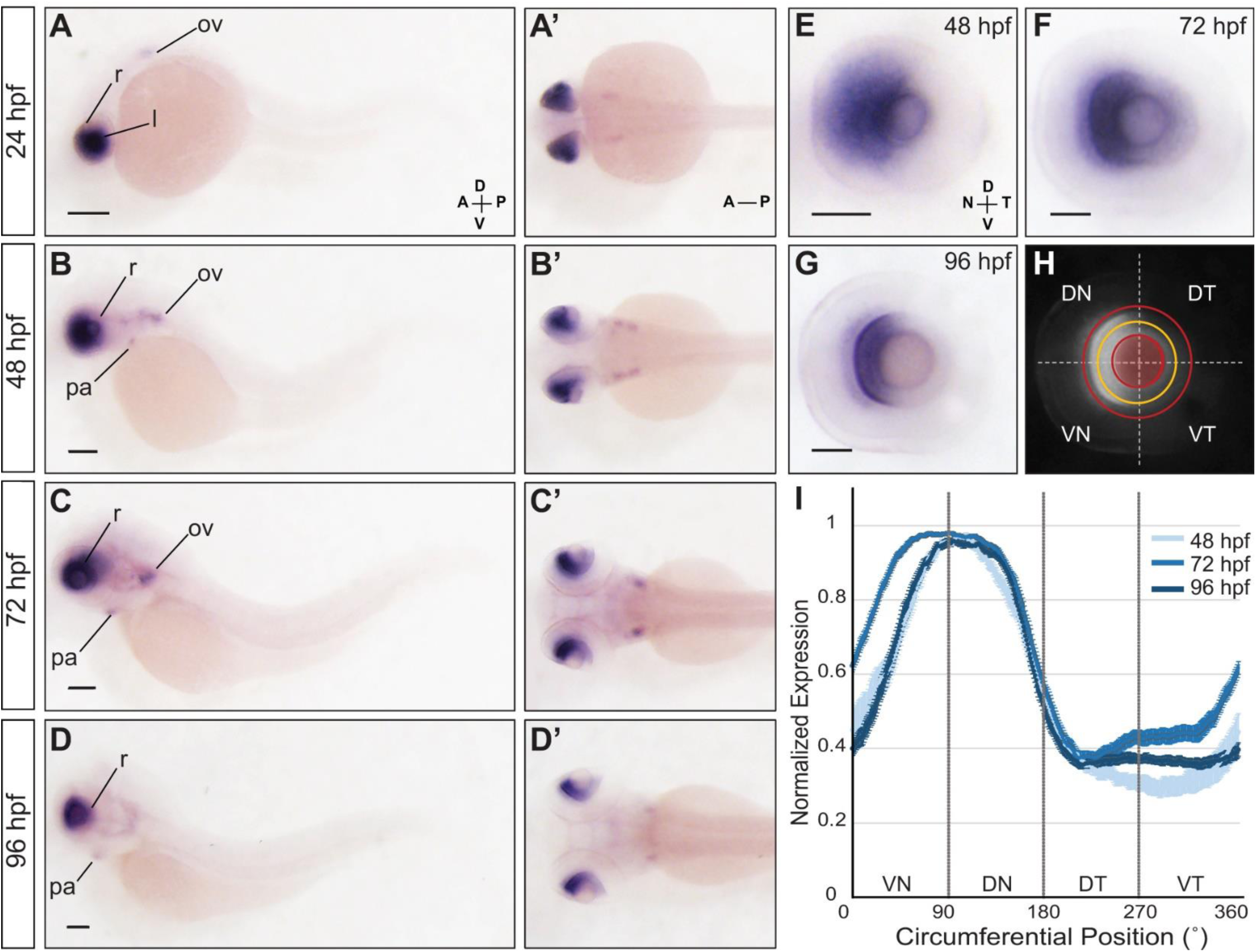
***Hmx1* is expressed in the nasal RGC layer throughout development**. Lateral (**A-D**) and dorsal (**A’-D’**) views of whole embryos stained for *hmx1* by ISH. (**A, A’**) At 24 hpf, *hmx1* is strongly expressed in the anterior retina (r) and lens (l) and weakly detected in the otic vesicle (ov). (**B, B’**) At 48 hpf, *hmx1* expression remains strongly detected in the anterior retina and is also seen in the otic vesicle and pharyngeal arches (pa). (**C, C’**) *Hmx1* expression remains consistent at 72 hpf. (**D, D’**) By 96 hpf, *hmx1* remains strongly expressed in the anterior retina and is still detected in the otic vesicle, and to a lesser extent, in the pharyngeal arches. (**E-G**) Lateral views of eyes dissected from embryos stained for *hmx1* by ISH. (**E**) At 48 hpf, *hmx1* is specifically detected in the nasal half of the retina. (**F, G**) This regionalized expression becomes restricted to the anterior RGC layer, and to a lesser extent the anterior inner nuclear layer, at 72 and 96 hpf. A similar expression is observed for *hmx4* (Figure 1 - supplement figure 1). (**H, I**) Quantification of *hmx1* expression in the RGC layer. (**H**) Intensity profiles of *hmx1* expression are measured on inverted grayscale images along a line (yellow) drawn half-way between the lens and the RGC layer periphery (delineated with red lines). (**I**) Quantification of signal intensity along a clockwise 360° trajectory shows that *hmx1* expression is restricted to the anterior RGC layer during retinotectal development. Means ± SEM are shown. Scale bars: 200 µm in A-D’; 50 µm in E-G. (A: Anterior, P: posterior, D: dorsal, V: ventral, N: nasal, T: temporal, VN: ventro-nasal, DN: dorso-nasal, DT: dorso-temporal, VT: ventro-temporal).

### A distal *hmx1* enhancer drives selective expression in nasal progenitors and RGCs

The restricted expression of *hmx1* and *hmx4* in the nasal retina prompted us to search for potential enhancers regulating their expression. Transcriptional enhancers are cis-regulatory elements containing short DNA sequences bound by specific transcription factors. Their activity has been correlated with the enrichment of specific post-translational modification of histones, allowing the prediction of their position in the genome. Active enhancers are notably associated with the presence of histone H3 lysine 4 monomethylation (H3K4me1) and H3K27 acetylation (H3K27ac), while H3K4me3 is predictive of active promoters (Bonn et al., 2012; Heintzman et al., 2009; Heintzman et al., 2007; Rada-Iglesias et al., 2011). We thus analyzed the genomic tracks of H3K4me3, H3K4me1, and H3K27ac modifications previously generated by (Bogdanovic et al., 2012) to identify putative distal regulatory elements in the *hmx1/4* locus region. *Hmx1* and *hmx4* are tightly linked on chromosome 14 and are both composed of two exons and one intron (**Figure 2A**). We identified two regions upstream *hmx1* that were characterized by the genomic co-localization of H3K4me1 and H3K27ac marks at 24 (data not shown) and 48 hpf (**Figure 2A**). We delineated a first putative element, *hmx1-En1,* as a 7 kb region right upstream the *hmx1* gene, and a second putative element, *hmx1-En2*, as a 7 kb region located more distally. A third putative regulatory element of 1.8 kb, *hmx1-En3*, was located in-between *hmx1* and *hmx4* genes. We tested the enhancer activity of these potential elements by generating stable transgenic lines expressing EGFP targeted to the plasma membrane by the CAAX prenylation motif of Ras (Moriyoshi et al., 1996) under the control of each of these elements. While *hmx1-En3* did not exhibit any enhancer activity, *hmx1-E*n1 and *hxm1-En2* drove EGFPCAAX expression in specific and partially overlapping regions at 96 hpf. Both enhancers were active in the pharyngeal arches and lips, but only *hmx1-En2* drove EGFP expression in the nasal retina and lens (**Figure 2B-C’**). Since developmental enhancers can be found in evolutionary conserved regions (Irimia et al., 2012; Woolfe et al., 2005), we used a multi-species alignment (visualized in UCSC genome browser) to identify conserved domains within *hmx1-En2.* We delineated a 3 kb region, *hmx1-En2s*, that was conserved across teleosts and amphibians. However, that region did not exhibit any enhancer activity despite its location within *hmx1-En2*.

**Figure 2.**
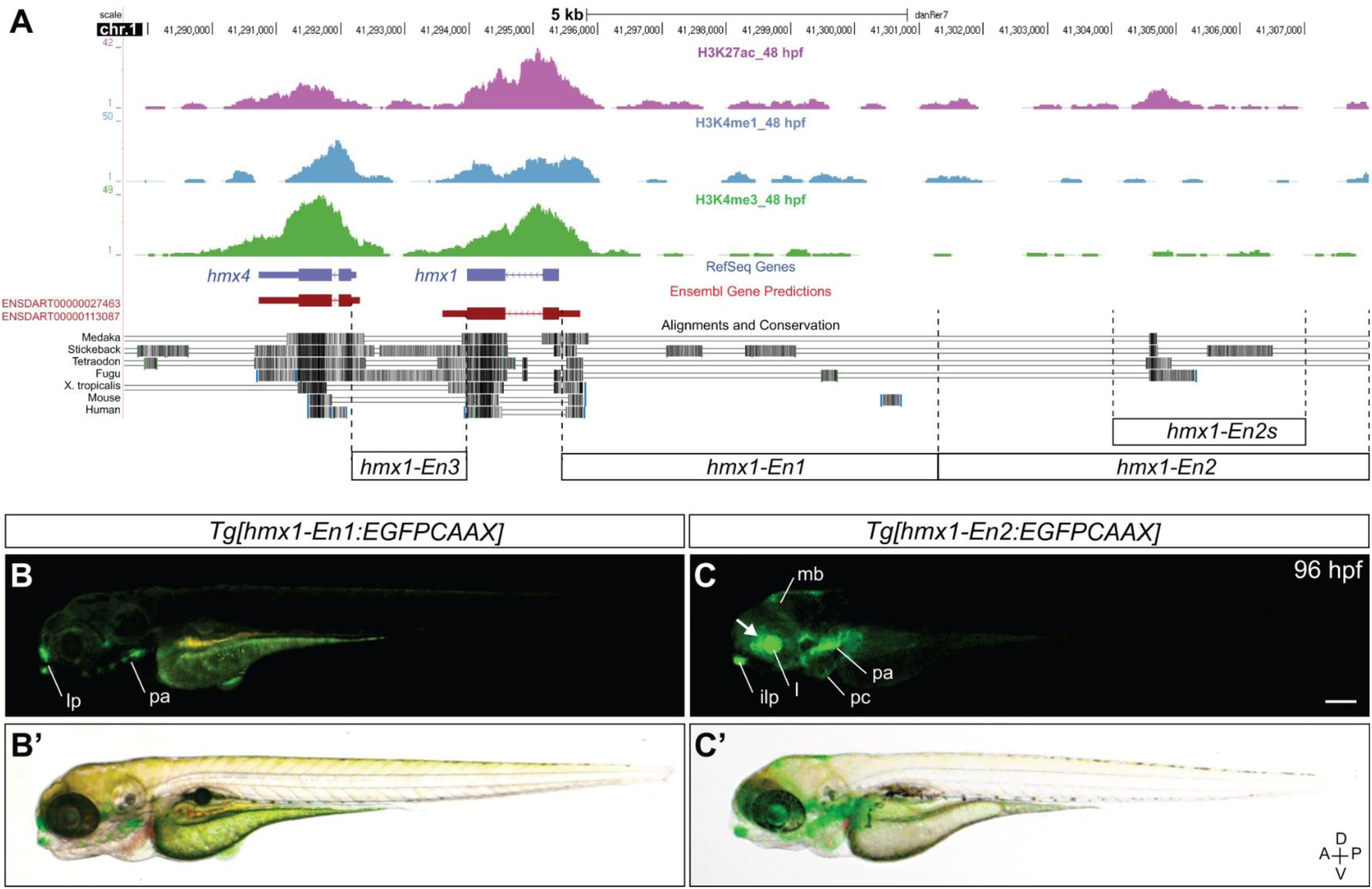
***Hmx1* enhancers recapitulate *hmx1* endogenous expression**. (**A**) Schematic representation of the *hmx4* and *hmx1* genomic locus on chromosome 1 (Zv9 assembly, UCSC Genome browser) (Kent et al., 2002). The distribution of H3K27ac, H3K4me1, and H3K4me3 modifications along 18 kb spanning the *hmx1*-*hmx4* locus at 48 hpf is shown (tracks from Bogdanovic et al., 2012). Four putative regulatory regions annotated as *hmx1-En1*, *hmx1-En2*, *hmx1-En2s* and *hmx1-En3* were tested for enhancer activity in stable transgenic embryos. (**B, B’**) The *hmx1-En1* enhancer drives EGFPCAAX expression in the pharyngeal arches (pa) and the lip (lp) region at 96 hpf. (**C, C’**) The *hmx1-En2* enhancer drives EGFPCAAX expression in the anterior retina (arrow), lens (l), midbrain (mb), pharyngeal arches (pa), inferior lip (ilp) and pericardic (pc) region at 96 hpf. Epifluorescence microscopy, scale bar: 200 µm.

As *hmx1-En2* was the only enhancer driving expression in the nasal retina at 96 hpf, we analyzed its activity throughout retinal development in more detail (**Figure 3**). At 24 hpf, before RGCs differentiate (Hu and Easter, 1999; Laessing and Stuermer, 1996; Schmitt and Dowling, 1999), EGFP expression was strongly detected in the nasal retina and lens of Tg[*hmx1-En2:EGFPCAAX*] transgenic embryos (**Figure 3A, A’).** This restricted expression was maintained at 48 and 96 hpf, with EGFP remaining selectively expressed in the nasal half of the retina at both stages (**Figure 3D-E’**). Interestingly, while *hmx1* transcripts were only detected in the RGC layer at 72 and 96 hpf (**Figure 1F, G**), EGFP remained visible in the entire nasal retina, likely because of its lasting stability in vivo. Like *hmx1* transcripts, EGFP was also found in several other structures including the pharyngeal arches at 48 and 96 hpf. It was also noticeably detected in the midbrain at 96 hpf (**Figure 3E, E’**). To determine whether *hmx1-En2* could drive transgene expression in nasal RGCs, we crossed our Tg[*hmx1-En2:EGFP*CAAX] transgenic line to Tg[*isl2b:TagRFP*] transgenic fish that express TagRFP under the control of the RGC-specific *isl2b* promoter (Pittman et al., 2008; Poulain and Chien, 2013). Confocal analysis of double transgenic embryos at 96 hpf revealed that EGFP partially overlapped with TagRFP in the nasal retina (**Figure 3G-H’**). Importantly, we could also detect EGFP in nasal retinal axons innervating the posterior half of the optic tectum in the midbrain (**Figure 3G, G’**), indicating that *hmx1-En2* is effective in driving selective expression in nasal RGCs at late stages of retinotectal development.

**Figure 3.**
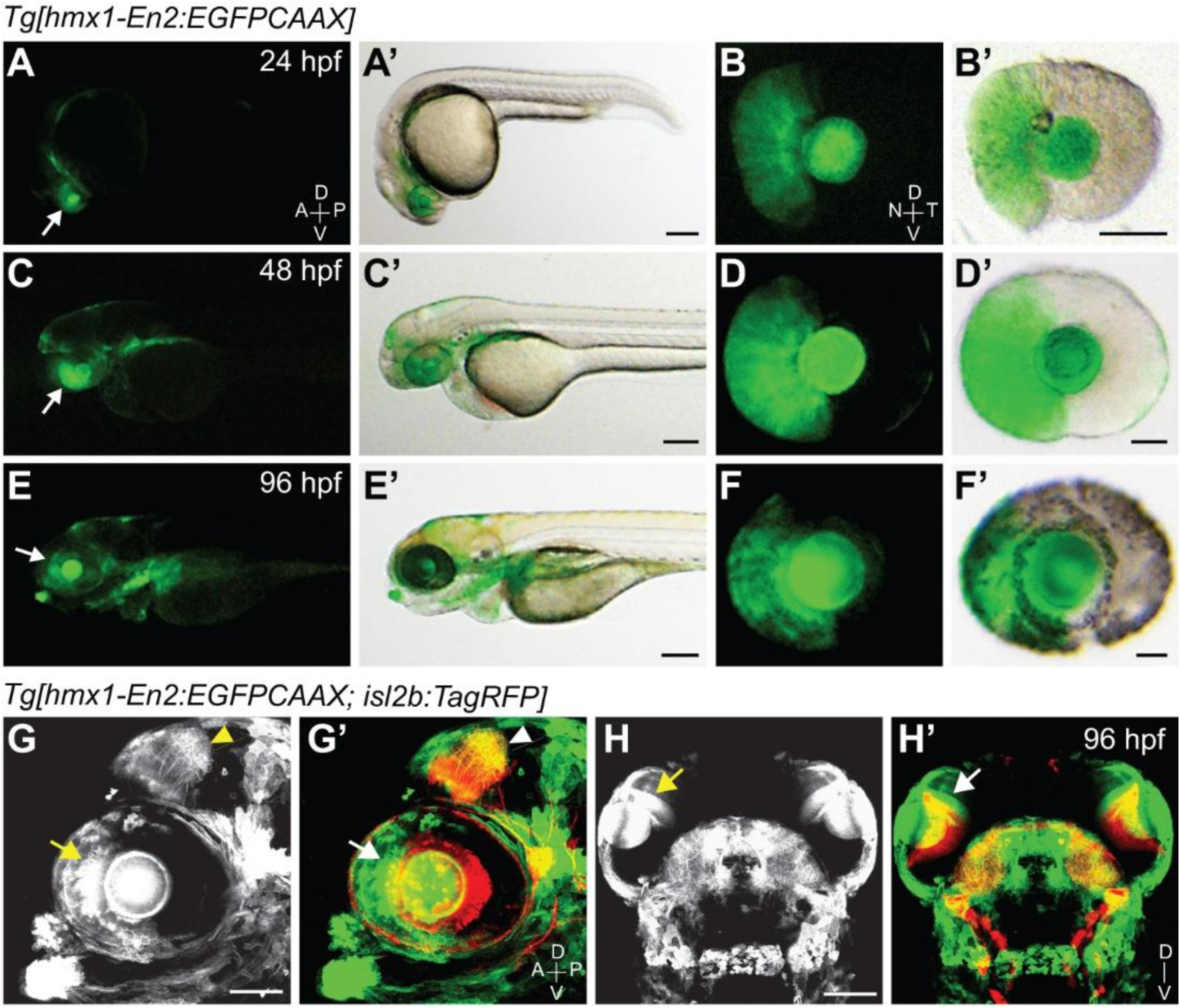
**The Hmx1-En2 enhancer drives expression in the anterior retina throughout development**. (**A-F’**) EGFPCAAX expression in [*hmx1-En2:EGFPCAAX*] transgenic embryos (A,C,E) and dissected eyes (B,D,F) at 24, 48 and 96 hpf. Fluorescence is detected in the anterior half of the retina (arrows) and the lens at all stages. No expression is observed in the posterior retina. Epifluorescence microscopy, scale bar: 200 µm in embryo pictures, 50 µm in eye pictures. (**G-H’**) Lateral (G, G’) and dorsal (H, H’) views of a double transgenic embryo expressing EGFPCAAX driven by the *hmx1-En2* enhancer and TagRFP driven by the RGC-specific *isl2b* promoter at 96 hpf. EGFPCAAX is observed in nasal RGCs (arrows) and nasal retinal axons projecting to the posterior tectum (arrowheads). Confocal microscopy, scale bar: 100 µm.

### *Hmx1:cre*-mediated recombination of an *RGC:colorswitch* reporter enables the visualization of the antero-posterior retinotopic map in vivo

Since the entire nasal retina and several brain structures beside the optic tectum were labeled in Tg[*hmx1-En2:EGFPCAAX*] transgenic larvae, we next sought to generate a stable transgenic line that would allow the direct visualization of the antero-posterior retinotectal map in vivo. The *Cre/loxP* system has been employed extensively in zebrafish for conditional expression and lineage tracing analyses using single-insertion loxP cassettes generated through Tol2-mediated transgenesis (Kawakami, 2007; Mosimann et al., 2011; Yoshikawa et al., 2008). We thus took advantage of that system to restrict transgene expression to nasal or temporal RGCs only. We generated a Tg[*hmx1-En2:cre*] stable transgenic line that expresses *cre* in the nasal retina, and a Tg[*isl2b:loxP-TagRFPCAAX-loxP-EGFPCAAX*] stable reporter line that expresses a switch transgene in RGCs (hereafter referred to as Tg[*RGC:colorswitch*]). To ensure that the *RGC:colorswitch* reporter transgene has integrated in an optimal genomic location for Cre-dependent recombination and to eliminate any functional positional effect, we established three independent Tg[*RGC:colorswitch*] stable transgenic lines and tested their responsiveness to Cre by crossing them to Tg[*hsp70l:cre*] transgenic fish (data not shown). We selected the Tg[*RGC:colorswitch*] reporter line whose progeny showed complete change of fluorescence in all RGCs following heat shock at 24 hpf (data not shown). We then crossed that line to generate a Tg[*hmx1-En2:Cre; RGC:colorswitch*] double transgenic line, and analyzed double transgenic embryos by immunolabeling for EGFP and TagRFP at 4 dpf (**Figure 4**).

**Figure 4.**
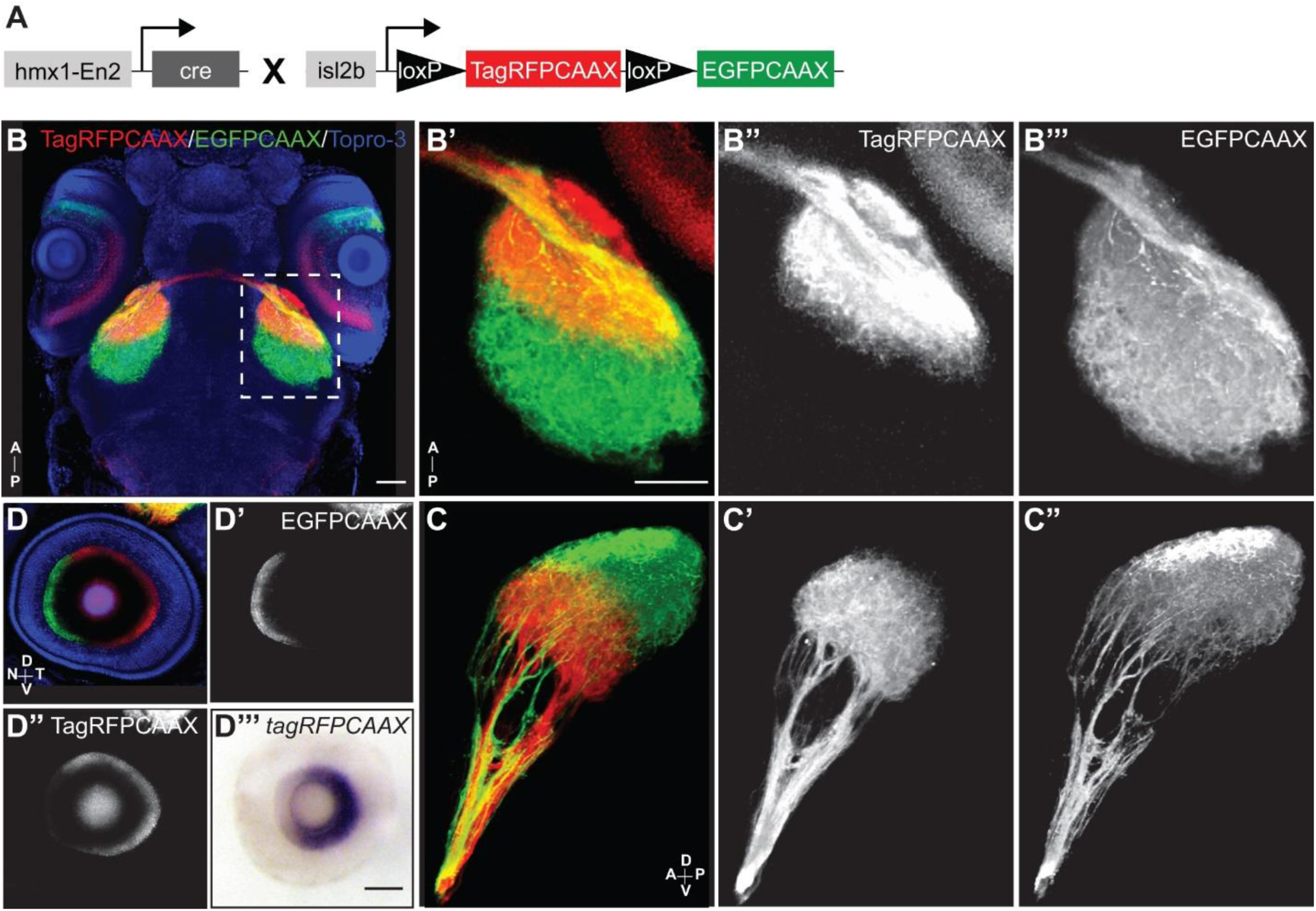
***Hmx1:cre*-mediated recombination of an *RGC:colorswitch* reporter enables the visualization of the antero-posterior retinotopic map in vivo.** (**A**) Schematic of the *hmx1-En2:cre* transgene and *isl2b:loxP-TagRFPCAAX-loxP-EGFPCAAX* (*RGC:colorswitch*) reporter expressed in double transgenic embryos. (**B**) Dorsal view of a double transgenic embryo immunostained for TagRFP and EGFP at 4 dpf. ToPro-3 was used as a nuclear counterstain to delineate the tectal neuropil. Corresponding 3D rendered movie of the antero-posterior retinotopic map is shown in Figure 4 - supplement movie 1. (**B’-B’’’**) 3D renderings of the tectum from a dorsal view. (**B”**) TagRFP-positive temporal retinal axons project specifically to the anterior half of the tectum. (**B’’’**) EGFP-positive nasal retinal axons project through the anterior tectum to the posterior tectum. (**C-C”**) 3D rendering of the optic tract and tectum from a lateral view. (**C**) Nasal and temporal retinal axons intermingle within the tract but project to distinct tectal areas. (**C’**) TagRFP-positive temporal axons project to the anterior half of the tectum. (**C”**) EGFP-positive nasal axons project to the posterior tectum. (**D-D”**) Eye of a double transgenic embryo immunostained for TagRFP and EGFP at 4 dpf. (**D’**) EGFP-positive RGCs are restricted to the nasal retina. (**D”**) TagRFP-positive RGCs are observed in the temporal half of the retina. (**D’’’**) Eye of a double transgenic embryo stained for *tagRFP* by ISH at 4 dpf. *TagRFP* expression remains restricted to the temporal retina. Confocal microscopy (**B-D”**), scale bar: 50 µm.

High resolution confocal imaging and 3D-rendering of double transgenic embryos revealed a bi-colored retinotectal map along the antero-posterior axis (**Figure 4, Figure 4-movie supplement 1**). We found that embryos had a bi-colored RGC layer in the retina, with nasal and temporal RGCs expressing EGFP and TagRFP, respectively (**Figure 4A, D-D”**). We confirmed by ISH that *tagRFP* was specifically expressed by temporal and not nasal RGCs at 4 dpf (**Figure 4D’’’**). We then analyzed the projection domains of nasal and temporal retinal axons at the tectum. After elongating together along both branches of the optic tract (**Figure 4C**), TagRFP-positive temporal retinal axons terminated in the anterior half of the tectum (**Figure 4B”, C’**) while GFP-positive nasal axons projected through the anterior tectum to reach the posterior tectum (**Figure 4B’’’, C”**). The sharp boundary between the nasal and temporal projection domains appeared to split the tectal neuropil into two equivalent halves (**Figure 4B’’’**). Thus, our observations indicate that *hmx1:cre*-mediated recombination can be used to drive selective transgene expression in nasal vs temporal RGCs. Our results also establish the Tg[*hmx1-En2:Cre; RGC:colorswitch*] transgenic line as the first genetic model allowing the direct visualization of retinotopic mapping in vivo throughout development.

### The antero-posterior retinotopic map is established at early developmental stages

We next analyzed retinotopic mapping in living larvae from 3 to 6 dpf (**Figure 5**). Like larvae fixed and immunolabeled at 4 dpf, living larvae had a bi-colored antero-posterior retinotectal map at all stages analyzed. To analyze retinotectal map development in a reproducible and unbiased manner, we established a consistent imaging and quantification method across embryos (**Figure 5 - figure supplement 1**). Confocal stacks of the retinotectal system taken from a dorsal view were consistently rotated along the x, y, and z axes to orient all embryos in a similar and comparable manner (**Figure 5A, A’; Figure 5 - figure supplement 1).** We then used maximal projections of rotated stacks to delineate several landmarks at the tectum and analyze the projection domains of nasal and temporal retinal axons (**Figure 5A”**). We used binarized maximal projections of TagRFP stacks to define the anterior-most boundary of the tectal neuropil (hereafter referred to as anterior tectal boundary) and the caudal boundary of the TagRFP-positive projection domain that we named “TagRFP Boundary”. We measured the distance (l) between the anterior tectal and TagRFP boundaries to determine the length of the TagRFP projection domain along the antero-posterior axis (**Figure 5 - figure supplement 1B-B”**). We used binarized maximal projections of EGFP stacks to set the posterior-most boundary of the tectal neuropil, and measured the total length of the tectum (L) as the distance between the anterior and posterior tectal boundaries. We also defined the Equator (E) as ½L, and used it to delineate the anterior (rostral to E) and posterior (caudal to E) halves of the tectum (**Figure 5 - figure supplement 1B-B”**).

**Figure 5.**
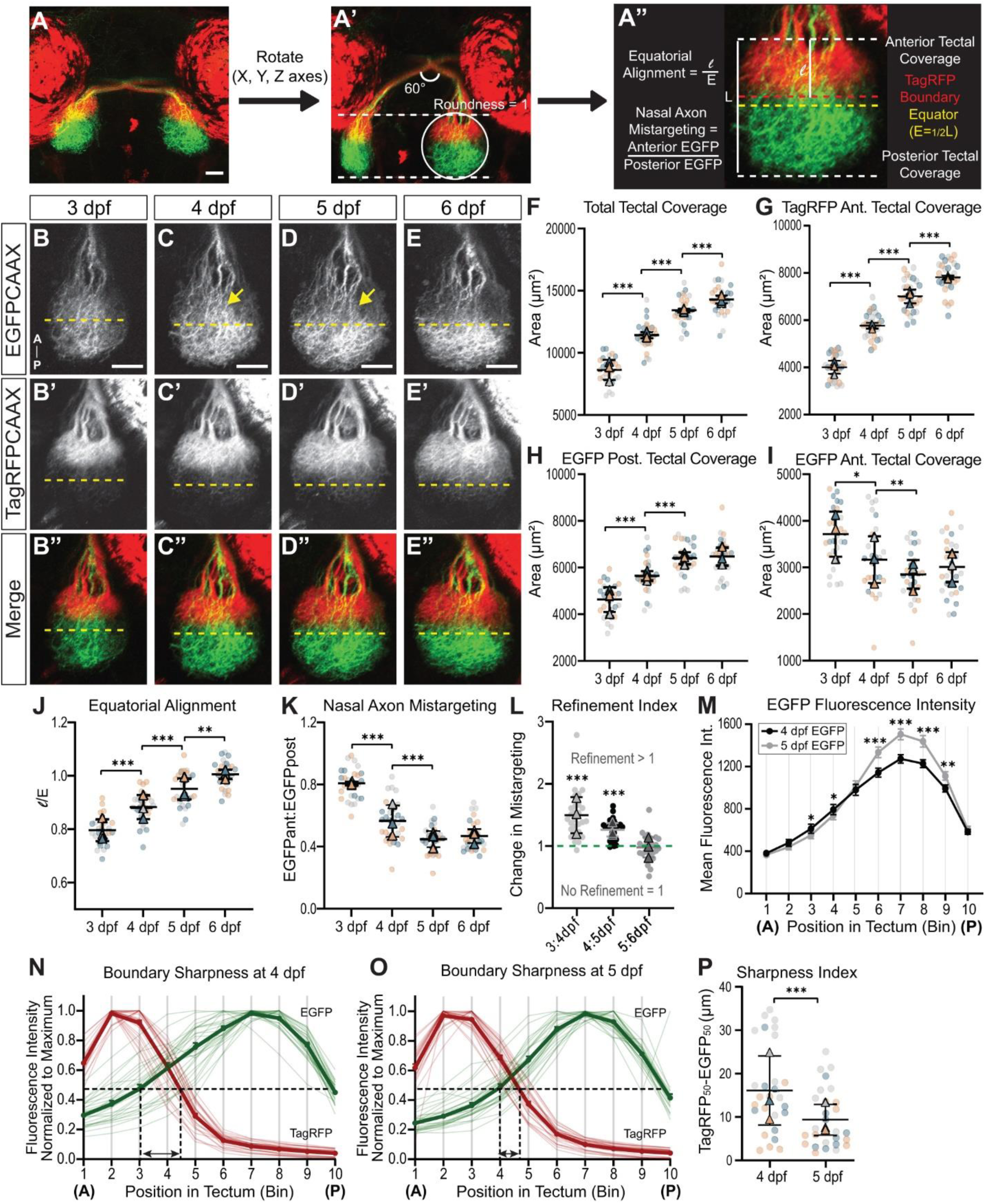
**Nasal retinal axons refine their tectal projection domain between 4 and 5 dpf.** (**A-A”**) Summarized quantification method to analyze retinotopic map development and refinement. A detailed description of quantification method is provided in Figure 5 - supplement figure 1. (**A-A’**) Z-series were opened in ImageJ and rotated so that both optic tracts intersect at an angle of 60°, and that the tectal roundness was equal to 1. (**A”**) After rotation, TagRFP and EGFP channels were separately maximum projected and thresholded for analysis of area coverage in the tectum. Four boundaries were defined to delineate the anterior and posterior halves of the tectum, with the anterior boundary corresponding to the rostral limit of the tectum (line where retinal axons enter the tectum), the posterior boundary corresponding to the caudal end of the tectum, the TagRFP boundary (red dashed line) corresponding to the caudal boundary of the TagRFP signal from temporal axons, and the Equator (E; yellow dashed line) corresponding to ½ of the total tectum length (L) measured from the rostral to the caudal tectal boundary. The tectal area rostral to the Equator was defined as the anterior half of the tectum, the tectal area caudal to the Equator, as the posterior half. (**B-E”**) Development of the antero-posterior retinotopic map from 3 to 6 dpf. (**B-E**) EGFP-positive nasal axons innervate the posterior half of the tectum by progressively refining their targeting domain. Between 4 and 5 dpf, nasal axons mis-targeting to the anterior half of the tectum seem to disappear (arrows). (**B’-E’**) TagRFP-positive temporal axons specifically and precisely target the anterior half of the tectum. (**B”-E”**) The antero-posterior topographic map is established fairly early on and maintained as retinal axons continue to innervate the tectum during development. As the tectum develops, the temporal retinal arborization field expands to fill the anterior half of the tectum, reaching the Equator position (yellow dashed lines) by 6 dpf. Confocal microscopy, scale bar: 50µm. (**F**) The total area of the tectum covered by TagRFP-positive and EGFP-positive axons significantly increases from 3 to 6 dpf. (**G**) The anterior area of the tectum covered by TagRFP temporal axons also steadily increases from 3 dpf and 6 dpf. (**H**) EGFP-positive nasal axons terminating in the posterior half of the tectum cover a significantly larger area between 3 and 4 dpf, and 4 and 5 dpf, before the area of coverage stabilizes between 5 and 6 dpf. (**I**) The anterior tectal area covered by EGFP nasal axons significantly decreases between 4 and 5 dpf, indicating a refinement of the nasal projection domain. (**J**) The Equatorial Alignment Index (l/E) corresponding to the ratio of the TagRFP coverage length (l) to the Equatorial length (E) steadily and significantly increases to a value around 1 from 3 to 6 dpf, indicating that the TagRFP boundary progressively shifts posteriorly until its position matches that of the Equator. (**K**) The Nasal Axon Mistargeting Index, defined as the ratio between the anterior and posterior tectal areas covered by EGFP-positive nasal axons, shows a significant decrease between 3 and 4 dpf as well as 4 and 5 dpf. (**L**) The Refinement Index that corresponds to the change in the Nasal Axon Mistargeting Index between two consecutive days is superior to one between 3 and 4 dpf and 4 and 5 dpf, indicating a refinement of nasal projections between these stages. It averages a value of 1 between 5 and 6 dpf, indicating that no refinement occurs during that time frame. (**M**) The mean fluorescence intensity of EGFP was measured in 10 bins of equal height along the length of the tectum at 4 and 5 dpf (a detailed description of quantification method is provided in Figure 5 - supplement figure 1). Bins 3 and 4 in the anterior half of the tectum show a significant decrease in signal between 4 and 5 dpf, while bins 6-9 in the posterior half of the tectum display a significant increase. (**N, O**) Normalized fluorescence intensities of EGFP and TagRFP were plotted along the antero-posterior axis of the tectum, and the distance from the anterior tectal boundary at which EGFP and TagRFP intensities reached 50% of their maximal value is marked by dashed lines. The distance between EGFP_50%_ and TagRFP_50%_ decreases between 4 and 5 dpf (double arrows), indicating that the boundary between EGFP and TagRFP projection domains becomes sharper over time. (**P**) The Boundary Sharpness Index corresponding to the absolute value of the distance between EGFP_50%_ and TagRFP_50%_ (double arrows in N and O) significantly decreases between 4 and 5 dpf. (**F-L, P**) Data represent mean ± SEM. n = 27 embryos. In each graph, three biological replicates representing independent experiments are color-coded in grey, teal and orange, respectively. Circle plots represent each data point, and triangle plots represent the averages of each biological replicate (Lord et al., 2020). Statistical Analysis: (F-K) repeated measures one-way ANOVA with Tukey’s posthoc test; (L) paired t-test compared to a control of 1 (1 representing no change); (M) paired t-test between 4 and 5 dpf within each bin; (P) paired t-test; *p < 0.05, ** p < 0.01, ***p < 0.001.

At 3 dpf, when the retinotectal map can first be visualized in fixed embryos (Stuermer, 1988; Burrill and Easter, 1995), we found that EGFP-positive nasal retinal axons had already elongated through the anterior half of the tectum to innervate the posterior half (**Figure 5B**). Some nasal axons also seemed to arborize in the anterior tectal half, just rostral to the Equator. On the other hand, TagRFP-positive temporal retinal axons projected specifically to the anterior tectum and were not observed in the posterior half (**Figure 5B’**). The TagRFP Boundary was rostral to the Equator at that stage (**Figure 5B’, B”**). From 4 to 6 dpf, the projection domain of TagRFP-positive temporal axons expanded within the anterior half of the tectum, thereby pushing the TagRFP Boundary towards the Equator (**Figure 5C’-E”**). Conversely, the projection domain of EGFP-positive nasal axons appeared denser and progressively more restricted to the posterior half of the tectum (**Figure 5C-E**). To better analyze the dynamics of retinotopic map formation, we quantified the tectal area covered by nasal and temporal axons over time (**Figure 5F-I, Figure 5-supplement figure 1B-B”**). Overall, the total tectal coverage (area of the tectum covered by nasal and temporal axons) significantly increased from 8,578.89 ± 202.23 µm^2^ at 3 dpf to 14,287.80 ± 243.06 µm^2^ at 6 dpf (p < 0.001), indicating a continuous growth and innervation of the tectum as development proceeds (**Figure 5F**). The area covered by TagRFP-positive temporal axons in the anterior half of tectum also steadily and significantly increased from 3,989.22 ± 99.55 µm^2^ at 3 dpf to 7,824.67 ± 130.46 µm^2^ at 6 dpf (p < 0.001) (**Figure 5G**), suggesting that the increasing innervation by temporal axons greatly contributes to the tectum growth. We further quantified the expansion of the area covered by temporal axons by measuring the position of the TagRFP Boundary in relation to the Equator (**Figure 5J**). The Equatorial Alignment Index corresponding to the ratio of the TagRFP coverage length (l) to the Equatorial length (E) gradually increased from 79.54 ± 1.0% at 3 dpf to 94.89 ± 1.2% at 5 dpf (p < 0.001). It eventually reached 100 ± 0.9% at 6 dpf, indicating that the TagRFP boundary progressively shifts posteriorly until its position matches that of the Equator. Finally, the area covered by EGFP-positive nasal axons in the posterior half of tectum also steadily and significantly increased from 4,589.66 ± 135.87 µm^2^ at 3 dpf to 6,381.07 ± 134.64 µm^2^ at 5 dpf (p < 0.001), but then remained stable from 5 to 6 dpf (**Figure 5H**). Thus, our results demonstrate that the antero-posterior retinotopic map is formed early on during development but remains dynamic, with the projection domains of both nasal and temporal axons expanding over time.

### Nasal retinal projections refine over time and generate a more precise map

In contrast to temporal axons that reach the anterior tectum immediately, nasal retinal axons must navigate through the anterior half of the tectum to reach their correct target in the posterior half. Interestingly, we noticed that some nasal axons seemed to arborize in the anterior tectal half just rostral to the Equator at 3 and 4 dpf (**Figure 5B, C**). However, these arborizations were not as clearly observed at 5 dpf, as shown by the apparent decrease in EGFP fluorescence intensity between 4 and 5 dpf (arrows in **Figure 5C, D**). We analyzed in more detail the tectal coverage of EGFP-positive nasal axons in the anterior half of the tectum. Strikingly, the area covered by nasal axons in the anterior tectum significantly decreased between 3 and 4 dpf (from 3,668.46 ± 119.45 µm^2^ to 3,175.32 ± 146.91 µm^2^, p < 0.001) and between 4 and 5 dpf (from 3,175.32 ± 146.91 µm^2^ to 2,840.33 ± 108.94 µm^2^, p < 0.001), but remained stable between 5 and 6 dpf (**Figure 5I**). While the values obtained at 6 dpf likely represent fluorescence from the nasal axonal bundles that have extended through the anterior tectum, the significant decreased in tectal coverage observed from 4 to 5 dpf suggests that nasal retinal projections in the anterior tectal half might refine during that time period. We thus calculated a Nasal Axon Mistargeting Index as the ratio between the anterior and posterior tectal areas covered by EGFP-positive nasal axons (**Figure 5K**). That index significantly decreased between 3 to 4 dpf (from 80.34 ± 1.93% to 56.76 ± 2.58%, p < 0.001) and between 4 to 5 dpf (from 56.76 ± 2.58% to 44.76 ± 1.72%, p < 0.001), but remained stable between 5 and 6 dpf (46.93 ± 1.83%). We also established a Refinement Index corresponding to the change in the Nasal Axon Mistargeting Index between two consecutive days (**Figure 5L**). The Refinement Index was superior to one between 3 and 4 dpf and 4 and 5 dpf (1.48 ± 0.07 and 1.27 ± 0.03, respectively), indicating a refinement of nasal retinal projections between these stages. In contrast, it averaged a value of 1 between 5 and 6 dpf (0.97 ± 0.04), indicating that no change or refinement occurred during that time period. Thus, our results indicate that nasal retinal axons refine and condense their projection domain to the posterior tectum from 3 to 5 dpf.

To determine the effect of that refinement on the retinotopic map, we decided to analyze the sharpness of the map at 4 and 5 dpf. We used sum projections of rotated stacks to measure the mean fluorescence intensity of EGFP and TagRFP in bins of equal height distributed along the antero-posterior axis of the tectum (**Figure 5 - figure supplement 1C-C”**). Interestingly, the mean EGFP intensity significantly decreased between 4 and 5 dpf in bins 3 and 4 in the anterior tectal half while it significantly increased in bins 6-9 in the posterior tectum (**Figure 5M**). We further analyzed the sharpness of the EGFP-TagRFP boundary by normalizing fluorescence intensities to their maximum values at 4 and 5 dpf, and plotting them along the antero-posterior axis of the tectum (**Figure 5N, O**). We then determined the distance from the anterior tectal boundary at which EGFP and TagRFP intensities reached 50% of their maximal value (dashed lines in **Figure 5N, O**). Interestingly, EGFP_50%_ shifted from an averaged position of 40.73 ± 13.16 µm at 4 dpf (anterior region of Bin 3) to 53.45 ± 9.92 µm (Bin 4) at 5 dpf. In contrast, TagRFP_50%_ kept a similar location between Bins 4 and 5 from 4 to 5 dpf (54.76 ± 5.22 µm and 59.76 ± 5.15 µm, respectively). We next calculated a Sharpness Index corresponding to the absolute value of the distance between EGFP_50%_ and TagRFP_50%_ (double arrows in **Figure 5N and O**). That index significantly decreased between 4 and 5 dpf (from 16.56 ± 1.95 µm to 9.63 ± 1.30 µm, p < 0.001) (**Figure 5P**), demonstrating that the boundary between nasal and temporal projection domains sharpens during that interval. Thus, our results demonstrate for the first time that the zebrafish retinotectal map sharpens and becomes more precise as nasal retinal projections refine and disappear from the anterior tectum.

### *Maco* mutants have retinotopic mapping and refinement defects

Previous studies have demonstrated a role for neural activity in regulating retinal axon arbor size and dynamics (Ben Fredj et al., 2010; Gnuegge et al., 2001; Hua et al., 2005; Kita et al., 2015; Munz et al., 2014; Ruthazer et al., 2003). We thus decided to test whether neural activity was required for the refinement of nasal axons in the anterior tectum by analyzing the retinotopic map of *macho* (*maco*) mutants at 4 and 5 dpf (Baier et al., 1996; Karlstrom et al., 1996; Trowe et al., 1996). *Maco* mutants harbor a mutation in the *pigk* gene that encodes a glycosylphosphatidylinositol (GPI)-anchor transamidase involved in GPI anchor synthesis and attachment to nascent proteins (Carmean et al., 2015; Ohishi et al., 2000). That mutation also causes a down-regulation of voltage-gated sodium channels leading to a lack of neural activity in peripheral sensory neurons and RGCs (Gnuegge et al., 2001; Granato et al., 1996; Ribera and Nusslein-Volhard, 1998). Both *maco* mutants and their wild-type (WT) siblings had a bi-colored antero-posterior retinotectal map at 4 and 5 dpf (**Figure 6A-D”**), with EGFP-positive nasal axons innervating the posterior tectal half (**Figure 6A-D**) and TagRFP-positive temporal axons, the anterior half (**Figure 6A’-D’**). However, we noticed differences in the anterior tectal coverage of nasal axons between WT and mutants. While nasal retinal projections seemed to disappear from the anterior tectum at 5 dpf in WT, they did not in *maco* mutants (**Figure 6C, D**). We quantified the tectal coverage of nasal and temporal axons at 4 and 5 dpf. While *maco* had a significantly smaller tectal neuropil than their WT siblings, the total tectal coverage significantly increased in both mutants and WT (**Figure 6E**). The area covered by EGFP-positive nasal axons in the posterior half of the tectum also significantly increased between 4 and 5 dpf in both *maco* and WT, although it was overall smaller in *maco* mutants (**Figure 6G**). In contrast, the area covered by nasal axons in the anterior half of the tectum significantly decreased from 4 to 5 dpf in WT but remained constant in *maco* mutants (**Figure 6H**). The Nasal Axon Mistargeting Index similarly decreased in WT (from 0.60 ± 0.02 at 4 dpf to 0.48 ± 0.02 at 5 dpf) but not in *maco* (0.60 ± 0.03 at both 4 and 5 dpf) (**Figure 6J**), demonstrating that the refinement of nasal projections does not occur in *maco* (**Figure 6H**). Lack of refinement was further confirmed by the Refinement Index that averaged 1 in *maco* (**Figure 6K**). Since we have shown that the refinement of nasal projections correlates with a sharpening of the retinotopic map (**Figure 5**), we next examined how the refinement defects observed in *maco* mutants might affect topographic mapping. Analysis of the EGFP-TagRFP boundary sharpness at 4 and 5 dpf revealed that the distance between EGFP_50%_ and TagRFP_50%_ did not change in *maco* mutants (**Figure 6 - figure supplement 1**). The Sharpness Index even slightly increased in *maco* (from 20.11 ± 1.97 µm at 4 dpf to 21.85 ± 2.05 µm at 5 dpf) instead of decreasing like in WT siblings (from 19.01 ± 1.75 µm to 15.13 ± 2.04 µm, p < 0.05) (**Figure 6L**). Thus, our results demonstrate that the lack of refinement of nasal retinal projections in *maco* prevents the sharpening of the retinotopic map over time.

**Figure 6.**
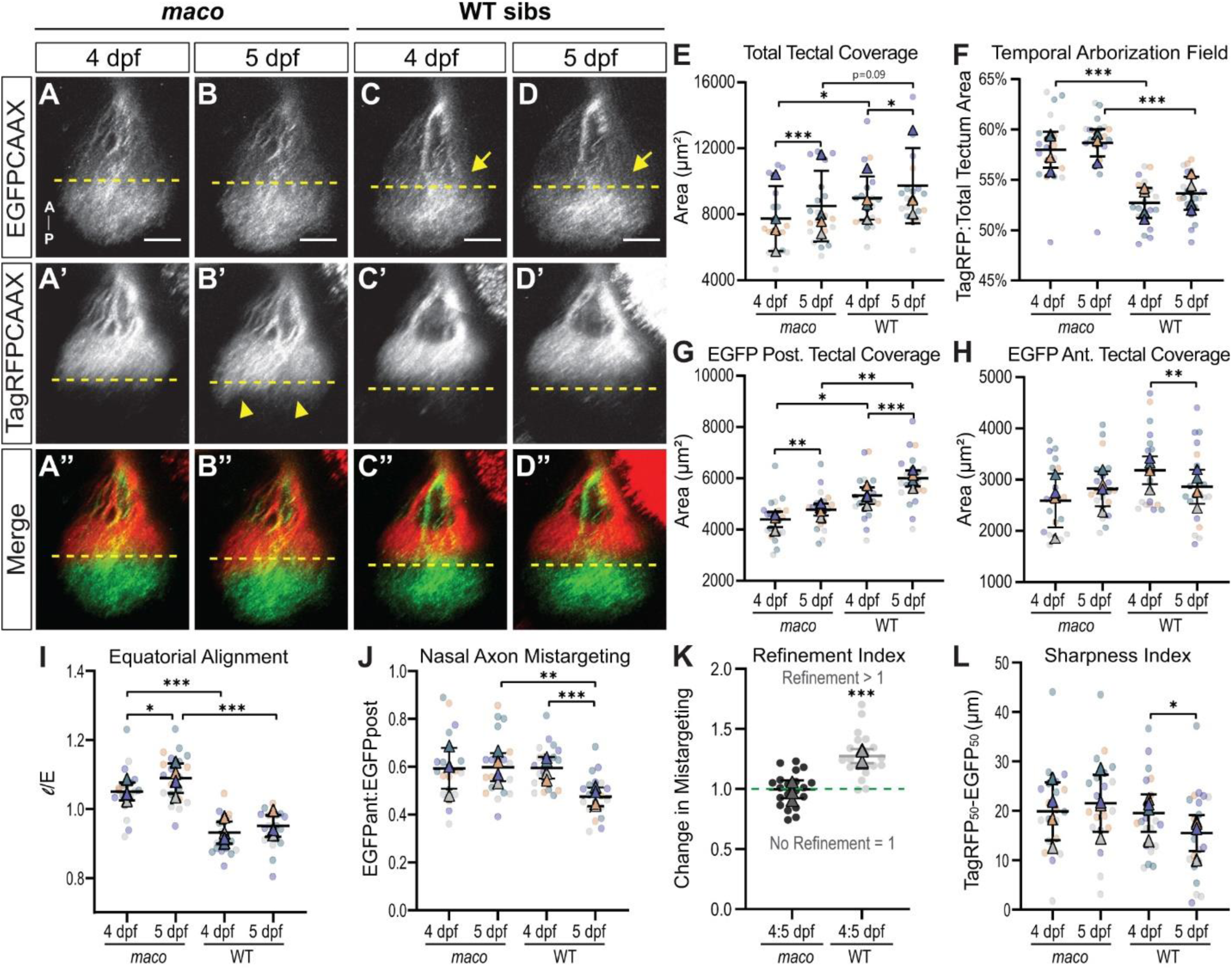
***Maco* mutants have retinotopic mapping and refinement defects**. (**A-D”**) Development of the antero-posterior retinotopic map in *maco* mutants (A-B”) and WT siblings (C-D”) from 4 to 5 dpf. (**A-D**) EGFP-positive nasal axons innervate the posterior half of the tectum in both *maco* and WT. The anterior tectal area covered by nasal retinal axons does not seem to change between 4 and 5 dpf in *maco* while it appears to decrease in WT siblings (arrows). (**A’-D’**) TagRFP-positive temporal axons specifically target the anterior half of the tectum in *maco* and WT siblings, but their projection domain expands caudally beyond the Equator (dashed line) in *maco* (arrowheads). Confocal microscopy, scale bar: 50µm. (**E**) Although *maco* have a significantly smaller tectum than WT at 4 and 5 dpf, the total area of the tectum covered by nasal and temporal axons significantly increases in both *maco* and WT between 4 and 5 dpf. (**F**) The Temporal Arborization Field index, defined as the ratio of the TagRFP area of coverage to the total tectal area, is significantly higher at 4 dpf and 5 dpf in *maco* compared to their WT sibling, indicating an expanded mapping of temporal retinal axons in *maco* mutants. (**G**) The area covered by EGFP-positive nasal axons in the posterior half of the tectum significantly increases from 4 to 5 dpf in both *maco* and WT. *Maco* have however a significantly smaller area of coverage in the posterior half of the tectum than WT at both stages. (**H**) The area covered by nasal axons in the anterior half of the tectum significantly decreases between 4 and 5 dpf in WT but not in *maco,* indicating an absence of refinement of the nasal projections in the mutant. (**I**) The Equatorial Alignment Index is greater than 1 at both 4 and 5 dpf in *maco,* indicating that the TagRFP boundary extends beyond the Equator in *maco* mutants. (**K**) The Nasal Axon Mistargeting Index decreases in WT but not in *maco* between 4 and 5 dpf, indicating a lack of refinement of the nasal projections in the mutant. (**L**) While the Refinement Index between 4 and 5 dpf is superior to 1 in WT, it averages 1 in *maco,* confirming the absence of refinement of the nasal projection domain in the mutant. (**M**) The Boundary Sharpness Index significantly decreases in WT but remains constant in *maco,* indicating that the boundary between EGFP and TagRFP projection domains does not refine over time in the mutant. Graphs representing the boundary sharpness at 4 and 5 dpf in maco are provided in Figure 6 - supplement figure 1. (**E-L**) Data represent mean ± SEM. n = 20 embryos per genotype. Four biological replicates representing independent experiments are color-coded in grey, teal, purple and orange, respectively. Statistical Analysis: (E-J, L) mixed-effects one-way ANOVA with Tukey’s posthoc test; (K) paired t-test compared to a control of 1 (1 representing no change); *p < 0.05, ** p < 0.01, ***p < 0.001.

In addition to the refinement defects affecting nasal axons, we unexpectedly observed subtle defects in the TagRFP-positive temporal projections in *maco*. We noticed that temporal axons arborized in a larger area in *maco* mutants than in WT siblings and expanded their area of coverage beyond the Equator (**Figure 6A’-D’**). We calculated a Temporal Arborization Field index as the ratio of the TagRFP area of coverage to the total tectal area, and found that it was indeed significantly higher at 4 and 5 dpf in *maco* compared to WT siblings (58.15 ± 0.77% vs 52.57 ± 0.51% at 4 dpf, and 58.80 ± 0.62% vs 53.44 ± 0.5% at 5 dpf) (**Figure 6F**). Moreover, the Equatorial Alignment Index exceeded 1 in *maco* at both 4 and 5 dpf (1.05 ± 0.2 and 1.09 ± 0.02, respectively) while remaining inferior to 1 in WT siblings at both stages (0.93 ± 0.01 and 0.94 ± 0.01, respectively) (**Figure 6I**). Our analysis thus revealed a slightly expanded mapping of temporal retinal axons in *maco* mutants. We could not observed any retinal patterning defects in *maco* (**Figure 6 - figure supplement 2**), suggesting that this topographic mapping defect results from local guidance errors at the tectum.

### Blocking neural activity in RGCs prevents the refinement of nasal retinal projections

The *pigk* mutation in *maco* not only affects other neurons besides RGCs but also likely prevents receptors other than voltage-gated sodium channels from being targeted to the plasma membrane. We thus decided to express the inward-rectifying potassium channel Kir2.1 (or a non-conducting form of Kir2.1, KirMUT, as a control) selectively in RGCs to test whether neuronal activity was required cell-autonomously for refining nasal retinal projections in the anterior tectum. We injected a *UAS:Kir2.1-2A-mKate2CAAX* or a *UAS:KirMUT-2A-mKate2CAAX* transgene in zygotes from an [*isl2b:gal4*] outcross, and analyzed whether transient expression of Kir2.1 blocked RGC activity by conducting a visually-mediated background adaptation (VBA) assay at 5 dpf (**Figure 7 - figure supplement 1**) (Neuhauss et al., 1999). The VBA is a physiological response dependent on RGC function during which zebrafish larvae adapt to changing levels of light by adjusting the distribution of melanin pigments (also known as melanosomes) in their skin (Kay et al., 2001). During normal VBA, melanosomes aggregate in response to bright illumination, giving larvae a pale appearance. We found that larvae expressing Kir2.1 in RGCs (thereafter referred to as Kir2.1 embryos) demonstrated two levels of dark, expanded pigmentation despite bright illumination (**Figure 7 - figure supplement 1B-C’**). Some Kir2.1 embryos retained dispersed melanosomes (embryos with “large melanophores”) in response to light, indicating a lack of VBA, while others had more restricted but yet abnormally expanded melanin (“smaller melanophores”), indicating a reduced VBA. In contrast, embryos expressing KirMut and uninjected embryos showed fully aggregated melanosomes in response to bright illumination (**Figure 7 - figure supplement 1D-E’**). We quantified the area covered by pigmented melanophores in these four groups and found that it was significantly larger in Kir2.1 embryos than in KirMUT or uninjected embryos (**Figure 7 - figure supplement 1F**). Transient expression of Kir2.1 in RGCs is thus sufficient to block neuronal activity in RGCs.

We then tested whether blocking RGC activity was sufficient to prevent the refinement of nasal retinal projections in the anterior tectal half. We selected embryos with high mKate2CAAX expression for our analysis (see Material and Methods for the definition of high mKate2 expression), and ensured by Western blot that high mKate2 expression correlated with high expression of Kir2.1 blocking RGC neural activity (**Figure 7 - figure supplement 1G**). As reported previously (Stuermer et al., 1990), blocking neuronal activity in RGCs did not prevent the overall formation of the retinotectal map (**Figure 7 - figure supplement 2A-F”**). The total tectal area covered by retinal axons significantly increased between 4 and 5 dpf in Kir2.1, KirMUT and uninjected embryos, with no significant difference between groups at either time point (**Figure 7A**). Similarly, no difference could be observed in the Temporal Arborization Field index (**Figure 7B**) or the Equatorial Alignment Index (**Figure 7C**). The area covered by EGFP-positive nasal axons in the posterior half of the tectum significantly increased between 4 and 5 dpf in all groups (**Figure 7D**). In contrast, the area covered by nasal axons in the anterior half of the tectum significantly increased from 4 to 5 dpf in Kir2.1 embryos (from 2,818.35 ± 130.98 µm^2^ to 3,140.43 ± 154.2 µm^2^, p < 0.05), while it decreased (from 2,933.79 ± 101.46 µm^2^ to 2,663.95 ± 89.66 µm^2^, p < 0.05) in uninjected embryos and tended to diminish (from 3,110.74 ± 133.37 µm^2^ to 2,950.5 ± 101.25 µm^2^) in KirMUT embryos. Consequently, the Nasal Axon Mistargeting Index remained constant in Kir2.1 embryos (from 0.61 ± 0.04 to 0.58 ± 0.03) instead of decreasing like in KirMut (from 0.63 ± 0.02 to 0.51 ± 0.02) and uninjected (from 0.53 ± 0.01 to 0.43 ± 0.05) control embryos (**Figure 7F**). Kir2.1 embryos had a Refinement Index averaging 1 (1.06 ± 0.03), demonstrating that their nasal projections did not refine in the anterior tectum between 4 and 5 dpf (**Figure 7G**). In contrast, KirMut and uninjected embryos had a Refinement Index of 1.24 ± 0.04 and 1.26 ± 0.04, respectively, indicating that refinement occurred during that time period. We finally examined the precision of the retinotopic map in Kir2.1, KirMUT and uninjected embryos (**Figure 7 - figure supplement 2G-L**). While the Sharpness Index significantly decreased between 4 and 5 dpf in KirMUT (from 26.14 ± 1.72 to 17.82 ± 1.36 µm, p < 0.01) and uninjected (from 18.21 ± 1.36 µm to 11.64 ± 1.44 µm, p < 0.05) embryos, it remained stable (from 23.76 ± 3.31 µm to 20.19 ± 2.23 µm) in Kir2.1 embryos (**Figure 7H**). Thus, our results demonstrate that selectively blocking neural activity in RGCs prevents the refinement of nasal retinal projections and the sharpening of the retinotopic map during development.

**Figure 7.**
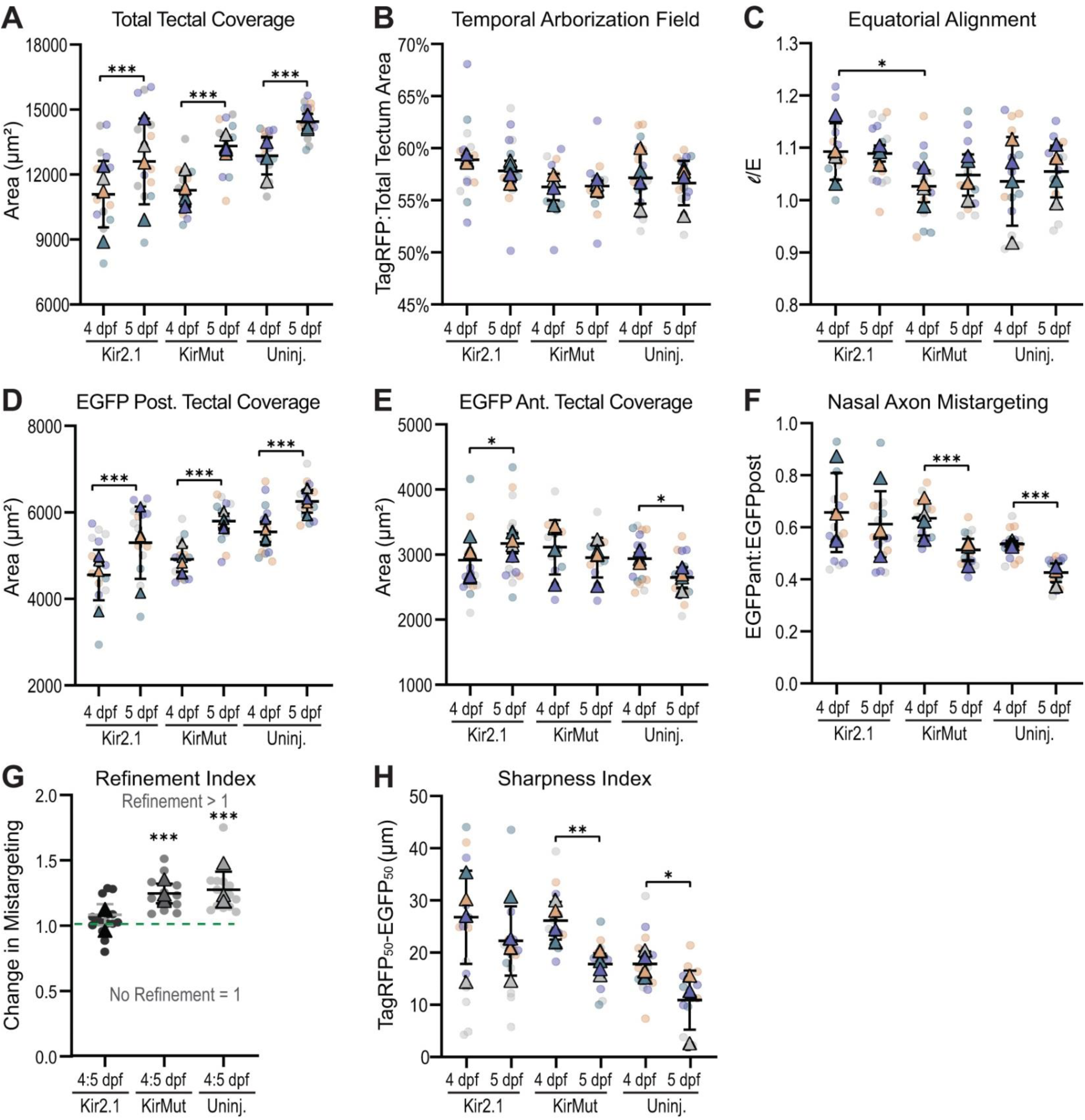
**Blocking neuronal activity in RGCs prevents the refinement of nasal projections**. (**A**) Like uninjected control embryos, embryos expressing Kir2.1 or KirMut in RGCs (thereafter referred to as Kir2.1 or KirMUT embryos) display an increase in total tectal coverage between 4 and 5 dpf. (**B**) The Temporal Arborization Field index is similar in Kir2.1, KirMUT and uninjected embryos at 4 and 5 dpf. (**C**) The Equatorial Alignment Index does not significantly differ between days or between experimental groups. (**D**) The area covered by EGFP-positive nasal axons in the posterior half of the tectum significantly increases between 4 and 5 dpf in Kir2.1 embryos but remains significantly smaller than that in the uninjected controls at both time points. (**E**) The anterior tectal area covered by EGFP-positive nasal axons slightly increases between 4 and 5 dpf in Kir2.1 embryos while it decreases in KirMut and uninjected embryos, suggesting a lack of refinement of the nasal projections in Kir2.1 embryos. (**F**) While the Nasal Axon Mis- targeting Index significantly decreases between 4 and 5 dpf in KirMut and uninjected embryos, it remains stable in Kir2.1 embryos, indicating a lack of refinement of the nasal projections. (**G**) While the Refinement Index between 4 and 5 dpf is greater than 1 in KirMut and uninjected embryos, it averages 1 in Kir2.1 embryos, confirming the absence of refinement of the nasal projection domain. (**H**) The Boundary Sharpness Index significantly decreases in KirMut and uninjected embryos but remains stable in Kir2.1 embryos, indicating that the boundary between the EGFP and TagRFP domains does not refine over time in embryos expressing Kir2.1 in RGCs. (**A-H**) Data represent mean ± SEM. n = 15 Kir2.1 embryos, 12 KirMut embryos, 15 uninjected embryos. Four biological replicates representing independent experiments are color-coded in grey, teal, purple and orange, respectively. Statistical Analysis: (A-F, H) mixed-effects one-way ANOVA with Tukey’s posthoc test; (G) paired t-test compared to a control of 1 (1 representing no change); *p < 0.05, ** p < 0.01, ***p < 0.001.

## DISCUSSION

Our understanding of topographic map development and maintenance has so far been limited by a lack of genetic models allowing the direct observation of maps over time. Here, we report the generation of a novel zebrafish transgenic line that, for the first time, allows for the unbiased and quantitative analysis of retinotopic map formation and refinement directly in vivo. Using live confocal imaging of transgenic larvae from 3 to 6 dpf, we show that the antero-posterior retinotopic map is formed at early developmental stages but remains dynamic as both retina and tectum grow, with the projection domains of nasal and temporal axons expanding over time. We further demonstrate that nasal retinal projections initially arborize in the anterior half of the tectum but progressively refine and condense their projection domain to the posterior tectum in an activity-dependent manner from 4 to 5 dpf. We finally show that the refinement of nasal retinal projections mediates the sharpening of the antero-posterior retinotopic map, and that both are prevented by blocking neuronal activity in RGCs.

In agreement with previous studies in zebrafish and other species (Boisset and Schorderet, 2012; Deitcher et al., 1994; Schulte and Cepko 2000; Stadler and Solursh, 1996; Yoshiura et al., 1998; Wang et al., 2000), our data reveal that the homeobox transcription factors *hmx1* and *hmx4* are expressed in a sharp nasal-high to temporal-low gradient in the retina throughout development (**Figure 1, Figure 1 - figure supplement 1**). The detection of *hmx1* and *hmx4* throughout the nasal retinal neuroepithelium at early stages indicates that both genes are expressed in proliferating neuroblasts and might regulate their positional identity and differentiation. Supporting that hypothesis, reduced *hmx1* expression has been shown to block retinal cell differentiation in zebrafish (Boisset and Schorderet, 2012; Schorderet et al., 2008) and cause microphthalmia in mouse (Munroe et al., 2009) and human (Gillepsie et al., 2015; Schorderet et al., 2008; Vaclavik et al., 2011). Although zebrafish embryos injected with *hmx1* morpholino oligonucleotides do not exhibit any eye patterning defect (Boisset and Schorderet, 2012), misexpression of *hmx1* or *hmx4* does alter the regional specification of the retina along the nasal-temporal axis in chick (Schulte and Cepko 2000; Takahasi et al., 2009), suggesting that *hmx1* and *hmx4* have redundant functions in teleosts. Interestingly, our analysis shows that the expression of both *hmx1* and *hmx4* becomes restricted to the nasal RGC and inner nuclear layers at later stages. We have also identified a distal regulatory element upstream *hmx1* and *hmx4* genes that drives expression in nasal RGCs at 4 dpf (**Figure 3**). Altogether, these data suggest that *hmx1* and *hmx4* are expressed by mature RGCs themselves at late stages of development. In support of that observation, recent studies using single cell profiling of the adult human retina have reported *hmx1* transcripts in RGCs as well as horizontal cells and Muller glia (Cowan et al., 2020; Lukowski et al., 2019; Voigt et al., 2019). Future studies examining the effects of manipulating *hmx1* expression directly in nasal or temporal RGCs will thus be of great interest to better understand the progression of the oculoauricular syndrome caused by *hmx1* mutations in human (Gillepsie et al., 2015; Schorderet et al., 2008; Vaclavik et al., 2011).

Using *hmx1:cre*-mediated recombination of an *RGC:colorswitch* reporter, we have generated a novel transgenic line that enables the unprecedented visualization of the antero-posterior retinotopic map in vivo. We were able to image and analyze for the first time retinotectal map development within the same embryo over successive days. This unparalleled temporal resolution allowed us to observe dynamic changes in retinotopic mapping that could not be seen previously in fixed embryos (**Figure 5**). We found that while the antero-posterior retinotopic map is established early on, it progressively shifts caudally as temporal axons expand their innervation of the anterior tectum. This caudal shift is accompanied by the progressive refinement of nasal projections that condense their projection domain to the posterior tectum. The consistent expression of *EGFP* and *tagRFP* transgenes in the nasal and temporal retina, respectively, allowed us to analyze in a reproducible manner multiple parameters of retinotopic mapping. Interestingly, these parameters did not show much variability across embryos with the same genetic background, indicating that retinotopic mapping is a robust and highly stereotyped process. That property allowed us to discover subtle phenotypes in mutants that were not detected in previous studies (**Figure 6**). We notably observed that temporal retinal axons arborize in a larger area and expand their coverage beyond the neuropil equator in *maco* mutants compared to WT siblings. This subtle mapping defect was not detected in embryos lacking neuronal activity in RGCs, suggesting that it might result from axon misguidance at the tectum. Interestingly, *maco* mutant harbor a mutation in Pigk, a component of the transamidase complex responsible for GPI anchor synthesis and attachment to nascent proteins (Carmean et al., 2015; Ohishi et al., 2000). Although it remains unknown which GPI-anchored proteins are impaired by Pigk deficiency, ephrin-As are good candidates. Ephrin-As and their receptors EphAs are indeed expressed in counter-gradients across the nasal-temporal axis in the retina and antero-posterior axis in the tectum and SC (Cheng et al., 1995; Connor et al., 1998; Drescher et al., 1995; Feldheim et al., 1998; Frizen et al., 1998; Higenell et al., 2011; Lambot et al., 2005; Marcus et al., 1996; Nakamoto et al., 1996; Scalia et al., 2009). Ephrin-A2 and ephrin-A5 are notably expressed at high levels in nasal RGCs and posterior tectum, while temporal RGCs express high levels of EphA receptors. Temporal axons are repelled by ephrin-As present in the posterior tectum and project more caudally in the SC of mice lacking different combination of ephrin-As (Feldheim et al., 2000; Nakamoto et al., 1996; Pfeiffenberger et al., 2006). Temporal axons also invade the target of nasal axons when ephrin-A5 is removed from both the retina and SC but not if ephrin-A5 is only lacking in the SC, indicating that nasal axons themselves participate in the repulsion of temporal axons from the caudal tectum (Suetterlin and Drescher, 2014). A dysregulation of ephrin-A targeting to the plasma membrane at the tectum and/or along nasal axons could thus explain the subtle phenotype we observed in *maco*.

While temporal axons expand their projection domain within the anterior tectum, we found that nasal retinal projections initially covering the caudal part of the anterior tectum refine and progressively condense their domain to the posterior half between 3 and 5 dpf. This refinement is unlikely to be caused by cell death in the retina, as apoptosis in the RGC layer peaks from 1.5 to 3 dpf before sharply decreasing from 4 to 6 dpf (Biehlmaier et al., 2001; Cole and Ross, 2001). Instead, it is likely driven by the dynamic rearrangement of axonal branching pattern as nasal axons extend caudally and arborize in their final zone in the posterior tectum. In xenopus, nasal axons retract their branches from the anterior tectum after initiating branches in both anterior and posterior tectal halves (O’Rourke and Fraser, 1990). While retinal axons in zebrafish were initially thought to elongate along straight trajectories and only arborize after reaching their target area (Kaethner and Stuermer, 1992), more recent studies using high-resolution time-lapse imaging have revealed a different mode of elongation where axons continuously extend and retract branches, and navigate by selective branch stabilization (Kita et al., 2015; Simpson et al., 2012). Once in their termination area, axonal arbors remain highly dynamic and frequently elongate and retract filopodia and branch tips, with only a small fraction of them being maintained in the mature arbor (Alsina et al., 2001; Ben Fredj et al., 2010; Campbell et al., 2007; Meyer and Smith, 2006; Munz et al., 2014; Ruthazer et al. 2006). A retraction of proximal branches or filopodia that have extended in inappropriate areas coupled with a constant remodeling of arborizing axons might thus redistribute the position of branches and cause arbors to shift caudally, leading to the progressive refinement of nasal projections we observed.

Our results demonstrate that the refinement of nasal projections requires neuronal activity in RGCs, as it does not occur in *maco* mutants lacking voltage-gated sodium channels or in embryos expressing the inward-rectifying potassium channel Kir2.1 in RGCs (**Figures 6 and 7**). Several studies have highlighted a role for neuronal activity in refining the size and morphology of terminal retinal arbors in the tectum and SC. Application of TTX increases branch dynamics and causes enlarged axonal arbors in xenopus and frog (Cohen-Cory, 1999; Reh and Constantine-Paton, 1985). TTX also prevents the refinement of retinal fibers that overshoot their termination zone in chick (Kobayashi et al., 1990). Similarly in the mouse, axons appear more diffuse and occupy a larger area in the SC after RGCs have been silenced by in utero electroporation of Kir2.1 (Benjumeda et al., 2013). Studies in zebrafish, however, have not detected any consistent changes in the size or morphology of terminal arbors after globally silencing RGCs. While the projection field of retinal axons appears more diffuse in *maco* mutants and embryos with silenced RGCs (Ben Fredj et al., 2010; Gnuegge et al., 2001; Trowe et al., 1996), the number of terminal branches or their total length does not appear to change at 5, 6 or 7 dpf, even though the formation of transient filopodia is increased (Ben Fredj et al., 2010; Gnuegge et al., 2001; Hua et al., 2005; Kaethner and Stuermer, 1994; Stuermer et al., 1990). What could explain these differences across species? First, the exact timing of analysis might have not allowed for the detection of subtle changes in arbor coverage. Analysis of retinal arbors in relation to their time of arrival at their termination zone has indeed revealed that silenced arbors continue to steadily and slowly expand their coverage after reaching their target instead of expanding and retracting like WT arbors (Kita et al., 2015). This continued expansion leads to a significant increase in coverage that was only detectable 14 hours after axons had reached their target. Alternatively, localized changes in a subset of branch tips could lead to a spatial rearrangement of axonal arbors without modifying the average branch number and length. Several studies in xenopus and the mouse have demonstrated that the pattern of neuronal activity has an instructive role in locally remodeling axonal arbors (Dhande et al., 2011; Munz et al., 2014; Rahman et al., 2020; Ruthazer et al., 2003). Activity is notably required for the elimination of branches whose firing pattern does not match that of their neighbors. An elegant study in xenopus has further demonstrated that the temporal order of retinal axon activity also regulates arbor dynamics and position (Hiramoto and Cline, 2014). Sequentially activating RGCs by a stimulus moving in an anterior-to-posterior (RGC activity propagating from the temporal to nasal retina), but not in a posterior-to anterior direction refines the topographic distribution of retinal axons along the antero-posterior axis. Axons stimulated earlier than convergent arbors (i. e. temporal axons) shift their arbors towards the anterior tectum, while axons stimulated later (i.e. nasal axons) shift their arbors posteriorly. In both cases, arbors modify their position by structurally rearranging their branches, with the spatial distribution of branch retraction dictating the direction of the shift. Nasal arbors that are stimulated after temporal arbors in forward-moving larvae lose branches located rostrally to their center of mass. Interestingly, spontaneous waves of retinal activity that preferentially initiate in the temporal retina and propagate to the anterior retina have been described in both the mouse and zebrafish (Ackman et al., 2012; Zhang et al., 2016). These directional waves are present as early as 2.5 dpf in zebrafish, suggesting that they might drive similar structural rearrangements of retinal arbors as those observed in the tadpole.

By promoting structural rearrangements of axonal arbors, spatiotemporal patterns of retinal activity drive the sharpening of visual circuits (Burbridge et al., 2014, Cang et al., 2005; Chandrasekaran et al., 2005; Stellwagen and Shatz, 2002; Xu et al., 2015; Xu et al., 2016; Zhang et al., 2011). Our data similarly show that blocking the refinement of nasal projections prevents the sharpening of the antero-posterior retinotopic map. While a lack of neuronal activity was known to cause less defined projection fields in zebrafish (Ben Fredj et al., 2010; Gnuegge et al., 2001), our study provides the first unbiased and precise quantification of topographic mapping over time in vivo, both in WT and after silencing RGCs. By analyzing the fluorescence intensity of nasal and temporal projection domains along the antero-posterior axis of the tectum, we were able to detect a gradual shift of nasal projections over time (reflected by a slope of fluorescence intensity becoming steeper) that is absent when RGCs were silenced. Overall, our observations and results from other studies suggest that the mechanisms of retinotopic map formation and refinement in teleosts and other species are more similar than previously thought. Retinal projections in mammals and chick eliminate misplaced branches in an activity-dependent manner after initially over-shooting their termination zone. Similarly in zebrafish, nasal retinal projections initially arborizing in anterior areas refine their projection domains in an activity-dependent manner. While increasing evidence indicate that both competitive and stabilizing interactions between axons play an important role in re-shaping arbors (Gosse et al., 2008; Hua et al., 2005; Louail et al., 2020; Rahman et al., 2020), how axons communicate and influence each other for refining their projection domains remains an important question. By enabling the selective manipulation of nasal and temporal RGCs, our new genetic model will provide new strategies for analyzing the molecular mechanisms by which axon-axon interactions contribute to precise retinotopic mapping in vertebrates.

## MATERIAL AND METHODS

### Zebrafish Husbandry and Maintenance

All experiments and procedures were approved by the Institutional Animal Care and Use Committee of the University of South Carolina. Zebrafish WT, mutant and transgenic embryos were obtained from natural matings, raised at 28.5°C in E3 medium (5 mM NaCl, 0.17 mM KCl, 0.33 mM CaCl_2_, and 0.33 mM MgSO_4_) in the presence of 150 mM of 1-phenyl-2-thiourea (PTU) (Millipore Sigma, Burlington, MA) to prevent pigment formation, and staged by age and morphology (Kimmel et al., 1995). WT embryos were from the Tübingen or AB strains. *Maco ^tt261^* heterozygous fish (generous gift from Dr. A. Ribera, University of Colorado School of Medicine) were identified by high-resolution melting analysis (HRMA) (Parant et al., 2009) using the following primers: *maco-hrma-fw*: 5’- TTGTACCGGTGACGAACG-3’, *maco-hrma-rv*: 5’- AACAGAAGAAGGCATGAATACAC-3’. *Maco* mutant embryos were identified based on their lack of touch response at 48 hours post-fertilization (hpf) (Granato et al., 1996). Embryos were anaesthetized in tricaine-S (Western Chemicals, Ferndale, WA) before fixation or imaging. Zebrafish larvae and young fish were nurtured using rotifer suspension and dry food (Gemma 75 and 150, Skretting USA, Westbrook, ME). Adult fish were fed with dry food (Gemma 300, Skretting USA).

### Cloning of *hmx* cDNAs and putative enhancers

For cloning *hmx1* and *hmx4* cDNAs, zebrafish mRNA was isolated from embryos at 24 and 48 hpf, respectively, using Trizol and the RNeasy mini kit (Qiagen, Hilden, Germany), and cDNA was prepared from RNA using the SuperScriptIII First-Strand Synthesis system (Invitrogen, Carlsbad, CA). *Hmx1* and *hmx4* cDNAs were amplified using the following full length primers: *hmx1-fw*: 5’-ATGCATGAAAAAAGCCAG- CAACAGC-3’, *hmx1-rv:* 5’-TCAGACAAGGCCTGTCATCTGC-3’, *hmx4-fw:* 5’-ATCTAACGGAGAATATGAG-CAAGGAG-3’, *hmx4-rv*: 5’-TCATATATCTCCATCAAACAGGCTGAAATAC-3’. Amplicons were subcloned into PCRII-TOPO (Invitrogen) and sequenced to verify gene identity and confirm sequence orientation. *Hmx1* putative enhancer elements were amplified by PCR from total genomic DNA using the LA Taq PCR kit v2.1 (TaKaRa, Mountain View, CA) and the following primers: *hmx1-En1-fw:* 5’-ACCGCACCACTAAA- GAGTCACAG-3’, *hmx1-En1-rv:* 5’-GGGTGATACGTGAATACCTCTAAGCA-3’, *hmx1-En2-fw:* 5’- GAGGGTGCCAGATGGAGATACAC-3’, *hmx1-En2-rv:* 5’-ACTGGCTCTGCTATGCTTCTGTTTC-3’, *hmx1-En2s-fw:* 5-GAACGGTACCGAACCGTCTATTAAAAGATTACACTAC-3’ (KpnI restriction site added in primer), *hmx1- En2s-rv:* 5’-GAACGGATCCAATAAACAAGGGACTAATAATTCAAGG-3’ (BamHI restriction site added in primer), *hmx1-En3-fw:* 5’-GAACGGTACCTCTTTGGAGACTGGCTGAACTGAC-3’ (KpnI restriction site added in primer), *hmx1-En3-rv:* 5’-GAACGGATCCATTCTCCGTTAGATGCGGGTCC-3’ (BamHI restriction site added in primer). Amplicons were purified on gel using the Qiaquick gel extraction kit (Qiagen), subcloned into PCRII-TOPO (Invitrogen) and sequenced before being digested and ligated into the Gateway *p5E-MCS* entry vector (Kwan et al., 2007) using the following restriction endonucleases: XhoI / BamHI (for *hmx1- En1*), DraII (for *hmx1-En2*), and KpnI / BamHI (for *hmx1-En2s* and *hmx1-En3*).

### Generation of transgenesis vectors

All transgenesis constructs were generated using the Tol2kit Gateway cloning system (Kwan et al., 2007). To generate *hmx1* reporter constructs, *p5E-hmx1-En1*, *p5E-hmx1-En2*, *p5E-hmx1-En2s, p5E-hmx1-En3, pME-EGFPCAAX* and *p3E-polyAv2* were recombined into the *pDestTol2pA2* destination vector using a Gateway Multisite LR reaction (Kwan et al., 2007) (Invitrogen). *p5E-hmx1-En2*, *pME-iCre* and *p3E- polyAv2* were recombined into *pDestTol2CR3* (*pDestTol2pA3* with *myl7:TagRFP* transgenesis marker) to generate the *hmx1-En2:iCre* construct. The sequence encoding *loxP-TagRFPCAAX-polyA-loxP* was amplified by PCR, purified on gel using the Qiaquick gel extraction kit (Qiagen), and recombined into the pDONR221 destination vector to generate the *pME-loxP-TagRFPCAAX-polyA-loxP* entry vector. A modified *p5E-isl2b-gata2a* entry clone encoding a 7.6 kb genomic fragment upstream of the *isl2b* start codon fused to the 1 kb promoter of *gata2a* (Ben Fredj et al., 2010; Pittman et al., 2008), *pME-loxP-TagRF- PCAAX-polyA-loxP, and p3E-EGFPCAAX-polyA* were recombined into the *pDestTol2pA2* destination vector to generate the *isl2b:loxP-TagRFPCAAX-loxP-EGFPCAAX* (*RGC:colorswitch*) reporter construct. *p5E- isl2b-gata2a*, *pME-gal4VP16* (Kwan et al., 2007), and *p3E-polyAv2* were recombined into *pDestTol2CG2* to generate the *isl2b:gal4* construct. The sequence encoding *Kir2.1-2A-mKateCAAX* flanked by FseI and DraI restriction sites was amplified by PCR using *pME-Kir2.1-2A-mGFP* (generous gift from Dr. J. Fetcho, Cornell University; Kishore and Fetcho, 2013) and *p3E-mKate2CAAX-polyA* as templates, purified on gel and recombined into pDONR221 to generate the *pME-Kir2.1-2A-mKateCAAX* entry vector. A non-conducting version of Kir2.1, KirMUT, was constructed similarly to generate the *pME-KirMUT-2A-mKate- CAAX* entry vector. *p5E-UAS* (Kwan et al., 2007), *pME-Kir2.1-2A-mKateCAAX* (or *pME-KirMUT-2A-mKate- CAAX),* and *p3E-polyAv2* were recombined into *pDestTol2pA2* to generate the *UAS:Kir2.1/KirMUT-2A- mKate2CAAX* construct.

### Generation of stable transgenic lines

Stable transgenic lines were generated using the Tol2 transposon method as described previously (Kawakami et al., 2000). 10 to 40 pg of purified DNA (*pTol2pA2-hmx1-En1:EGFPCAAX, pTol2pA2-hmx1- En2:EGFPCAAX, pTol2pCR3-hmx1-En2:iCre, RGC:colorswitch, pTol2CG2-isl2b:gal4)* were co-injected with 25 pg of synthetic mRNA encoding Tol2 transposase at one-cell stage, and injected embryos with transient expression of transgenes were raised up to adulthood as F0 generation. F0 fish were then out- crossed to WT to screen for positive F1 embryos expressing the transgenes. Expression of EGFP and TagRFP driven in the heart by the *myl7* promoter was used to identify *isl2b:gal4* and *hmx1-En2:iCre* carriers, respectively. Transgenic F1 carriers were subsequently out-crossed to WT to generate stable lines with a single-copy insertion.

### Transient expression of Kir2.1 and Kir2.1MUT

For transiently expressing Kir2.1 or KirMUT together with mKate2CAAX in RGCs, 40 pg of *UAS:Kir2.1-2A- mKate2CAAX* or *UAS:KirMUT-2A-mKate2CAAX* were co-injected with Tol2 transposase mRNA at one cell stage in zygotes obtained from a cross between [*isl2b:gal4*] and [*hmx1:iCre; RGC colorswitch*] stable transgenic lines. Triple transgenic embryos identified by the expression of both TagRFP and EGFP in the heart and RGCs were selected for imaging and quantification. As reported in other systems (Eckardt et al., 2004; Luo et al., 2020; Schmidt-Supprian and Rajewsky 2007), about 5% of [*hmx1:iCre; RGC colorswitch*] double transgenic embryos showed aberrant Cre-mediated recombination that was first observed in the trigeminal at 24 hpf. These embryos were discarded from our analysis.

### Whole-mount In Situ Hybridization

For making ISH probes, cDNA templates cloned into pCRII-TOPO were amplified by PCR using M13fw and M13rv primers and purified on gel. In *vitro* transcription of digoxigenin-labeled probes was performed using the DIG RNA Labeling Kit (Millipore Sigma) according to manufacturer’s instructions. Embryos were dechorionated at the appropriate developmental stages and fixed in 4% paraformaldehyde in phosphate buffered saline (pH 7.4) for 2 hours at room temperature and overnight at 4°C. Whole-mount in situ hybridization was performed as previously described (Thisse and Thisse, 2008). After staining, embryos were cleared in 80% glycerol for imaging. Sense probes were used as controls and did not reveal any staining. Images were acquired using an Olympus SZX16 stereomicroscope equipped with an Olympus DP80 dual color camera and Cellsens standard software. Digital images were cropped and aligned using Adobe Illustrator.

### Quantification of retinal gene expression

Quantification of gene expression in the retina was carried out according to (Picker and Brand, 2005) with the following modifications. Eyes were dissected from embryos stained by ISH using a sharpened tungsten needle and imaged in 80% glycerol in a lateral view as described above. Images were imported into Fiji ImageJ analysis software (Schindelin et al., 2012; Schneider et al., 2012), transformed to 8-bit grayscale images, and inverted. An oval selection was applied half-way between the lens and the RGC layer periphery, and signal intensity was measured along a 360⁰ trajectory using the Oval Profile Plot plugin. Values were exported and analyzed in Microsoft Excel.

### Immunolabeling

Embryos were fixed in 4% PFA in Phosphate-buffered saline (PBS) for two hours at room temperature and then overnight at 4°C. Embryos were washed three times in PBT (PBS + 0.5% Triton X-100). Antigen retrieval was done in 150 mM Tris, pH 9 for 5 minutes at room temperature followed by 20 minutes at 70°C. Embryos were then permeabilized at room temperature for 15 minutes in water first, and then for 30 minutes in PBS with 1% Triton and 0.1% collagenase. Embryos were blocked for one hour at room temperature with blocking buffer (PBS with 0.5% Triton, 1% DMSO, 1% BSA, and 2% normal goat serum). Primary anti-EGFP (ab290, Abcam, Cambridge, UK) and anti-TagRFP (M204-3, MBL International, Woburn, MA) antibodies were applied at 1:500 dilution in PBT supplemented with 1% DMSO and 1% BSA overnight for 4°C. Embryos were washed three times in PBT. Secondary Alexa Fluor 488 Goat anti-Rabbit (111-545-003, Jackson Immuno Research, West Grove, PA) and Alexa Fluor 594 Donkey anti-Mouse (715-585-150,Jackson Immuno Research) antibodies were diluted 1:500 in PBT with 1% DMSO and 1% BSA and applied together with TO-PRO-3 (T3605, Thermo Fisher, Waltham, MA) diluted 1:1000 overnight at 4°C. Embryos were washed five times in PBT and mounted in 1% ultrapure low melting point (LMP) agarose (16520050, Thermo Fisher) for confocal imaging.

### Confocal Microscopy

For time-course imaging of live embryos from 3 to 6 dpf, embryos were anesthetized in 0.015% tricaine-S and embedded dorsally in 1% LMP agarose in E3 medium + PTU in a lumox membrane-bottomed dish (Greiner Bio-one, Monroe, NC). Images were acquired on a Leica TCS SP8X laser-scanning confocal microscope equipped with LASX software, HyD detectors, and a 20X objective. Z-series of the entire retinotectal system were acquired at 512 × 512 pixel resolution with a zoom of 1 and Z-intervals of 1.5 µm. After imaging, embryos were kept individually in a 12-well dish and allowed to recover for 24 hours before being re-anesthetized and mounted dorsally for the next day of imaging. Maximal and sum intensity projections were compiled at each time point in ImageJ software.

For high resolution imaging of the retinotectal system at 4 dpf, immunolabeled embryos were mounted either laterally after removing the contralateral eye or dorsally in 1% LMP agarose and imaged as described above with a Z-interval of 1 µm. 3D reconstructions of the retinotectal system were generated using FluoRender (Wan et al., 2012; Wan et al., 2017).

### Quantification of Topographic Mapping at the Tectum

All quantitative analyses of topographic mapping were conducted using ImageJ software. For unbiased analysis, dorsal view Z-series were consistently rotated along the X, Y, and Z-axes using the TransformJ Plugin (Meijering et al., 2001), so that the left and right tecta were aligned horizontally, both optic tracts intersected at an angle of 60°, and the roundness of the right tectal neuropil was equal to 1. To analyze the area of coverage, rotated images were maximally projected and binarized using a threshold of 80 for TagRFP and of 75 for EGFP. Thresholds were chosen to best represent the raw images acquired. Using the TagRFP channel, we delineated the anterior tectal boundary as the anterior line where retinal axons enter the tectum, and the TagRFP posterior boundary. Using the EGFP channel, we delineated the posterior tectal boundary as the caudal-most border where retinal axons arborize at the tectum. We defined the Equator (E) as ½ the length of the tectum measured along the antero-posterior axis between the anterior and posterior tectal boundaries. We used the Analyze Particles tool in ImageJ to measure the tectal coverage (area) of the TagRFP-positive temporal axons, of the EGFP-positive nasal axons in the anterior half of the tectum (rostral to the Equator), and of the EGFP-positive nasal axons in the posterior half of the tectum (caudal to the Equator). We calculated The Total Tectal Coverage as the sum of the TagRFP and posterior-EGFP axonal coverages.

To further analyze retinotopic mapping along the antero-posterior axis, we defined an Equatorial Alignment Index as the ratio between the antero-posterior length (l) of the TagRFP coverage (measured between the anterior tectal boundary and the TagRFP boundary) and the Equator. We established a Nasal Axon Mistargeting Index as the ratio between the EGFP area of coverage in the anterior half of the tectum and the EGFP area of coverage in the posterior half of the tectum. We calculated a Refinement Index as the ratio change of the Nasal Axon Mistargeting Index between two consecutive days.

To quantify the sharpness of the boundary between the TagRFP and EGFP projection domains, rotated images were sum-projected for measuring the mean fluorescence intensity of the TagFRP and EGFP signals. We divided the antero-posterior length of the tectum into 10 bins of equal height on merged images using the Polygon Selection tool in ImageJ, and measured the mean fluorescence intensity of each channel in each bin using the Measurement function in ImageJ. Bin1 was defined as the anterior-most tectal bin and Bin10, as the posterior-most tectal bin. Intensity values were normalized to the maximum value for each channel and plotted along the antero-posterior axis. We determined the point at which fluorescence intensity reached 50% of its normalized maximum value for each channel, and defined the Sharpness Index as the absolute distance between the EGFP_50%_ and TagRFP_50%_ positions at 4 and 5 dpf. (See **Figure 5 - supplement figure 1** for a detailed illustration of our quantifications).

For analyzing retinotopic mapping in *maco* mutants, we imaged embryos from a *maco*/+ incross in blind conditions and genotyped embryos after imaging. Only mutants and WT siblings were kept for quantification. A few *maco* mutants developed periocular edema by 5 dpf and were discarded from our analysis.

For analyzing retinotopic mapping in embryos expressing Kir2.1 or KirMUT in RGCs, we imaged mKate2CAAX at the tectum of injected embryos (see previous section on transient expression of Kir2.1 transgene) at 4 dpf using an excitation wavelength shifted to 615 nm to prevent any bleed-through of the TagRFP signal. We measured the mean fluorescence intensity of mKate2CAAX over the total tectal area and selected embryos with an average intensity over 400 for our analysis (referred to as “high mKate2” embryos). Of note, mKate2CAAX could be visualized along the optic tracts in “high mKate2” embryos but not in “low mKate2” embryos with these imaging conditions.

### Visually-mediated Background Adaptation (VBA) Assay

Larvae were exposed to bright illumination from the bottom for 30 minutes to light-adapt and then imaged from a dorsal view at 5 dpf. Images were binarized using a threshold of 40, and pigmentation of larvae was assessed by measuring the area of melanophore coverage in a region caudal to the eyes and rostral to the medulla oblongata (See **Figure 7 – supplement 1**).

### Western Blotting

Embryos were anaesthetized at 5 dpf and lysed in RIPA buffer (Millipore Sigma). Protein concentrations were measured using a Pierce BCA Protein Assay Kit (Thermo Fisher), and 50 µg of denatured proteins were run on a 12% acrylamide gel and transferred to a 0.2 µm nitrocellulose membrane (BioRad, Hercules, CA). We blocked membranes in 5% Milk in TBST (Tris Buffer Solution + 0.1% Tween) and applied primary antibodies overnight at 4°C. Secondary antibodies were applied for 1.5 hours at room temperature, and proteins were detected using Amersham Western Blotting Detection Reagent (GE Healthcare, Chicago, IL). Western blot membranes were probed with mouse anti-RFP (RF5R Invitrogen, 1:500 dilution), rabbit anti-Kir2.1 (APC-026, Alomone Labs, Jerusalem, Israel, 1:500 dilution), and rabbit anti-GAPDH (14C10, Cell Signaling Technology, Danvers, MA, 1:2000 dilution). Secondary HRP-conjugated horse-anti-mouse (7076S, Cell Signaling Technology) and goat-anti-rabbit antibodies (7074S, Cell Signaling Technology) were used at a 1:2000 dilution.

### Statistical Analysis

All statistical analyses were performed with GraphPad Prism 9 software (GraphPad, San Diego, CA). We define biological replicates as individual larvae from a mixed clutch born from pairings of at least two males and two females. Each experiment was repeated at least three times under similar experimental conditions. Sample size was decided was based on the low variability detected in pilot studies. Data are presented as mean ± SEM. For better communicating variability across samples and experimental reproducibility, graphs are represented as “SuperPlots” (Lord et al., 2020), in which biological replicates representing independent experiments are color-coded, with circles representing individual embryos tested and triangles representing averages. To compare groups with repeated measures over time, we performed repeated measures one-way ANOVA followed by Tukey’s multiple comparisons post-hoc test. To compare data with both “within-subjects” and “between-subjects” factors (i.e. *maco* and Kir2.1 data), we performed mixed effects one-way ANOVA followed by Tukey’s multiple comparisons post-hoc test. The number of larvae analyzed and the statistical methods used to compare groups are described in details in each figure legend.

## ACKNOWLEDGEMENTS

This work was supported by the National Institutes of Health/ National Institute of Neurological Disorders and Stroke (R01NS109197 to F.E.P.), the University of South Carolina SmartState Center in Childhood Neurotherapeutics (to F.E.P.) and an Aspire I grant from the Office of the Vice President for Research at the University of South Carolina (to F.E.P). We would like to thank Dr. Angie Ribera (University of Colorado School of Medicine) for providing the *maco* mutant line and Dr. Joe Fetcho (Cornell University) for giving us constructs encoding Kir2.1 and KirMUT. We also thank expert help of Dr. Edsel Pena (University of South Carolina) for assistance with statistical analyses, and Dr. Cory Weaver for his helpful comments on mapping analysis. We finally thank Quill Thomas for technical assistance and fish husbandry.

## FIGURE LEGENDS

**Figure 1 - supplement figure 1.**
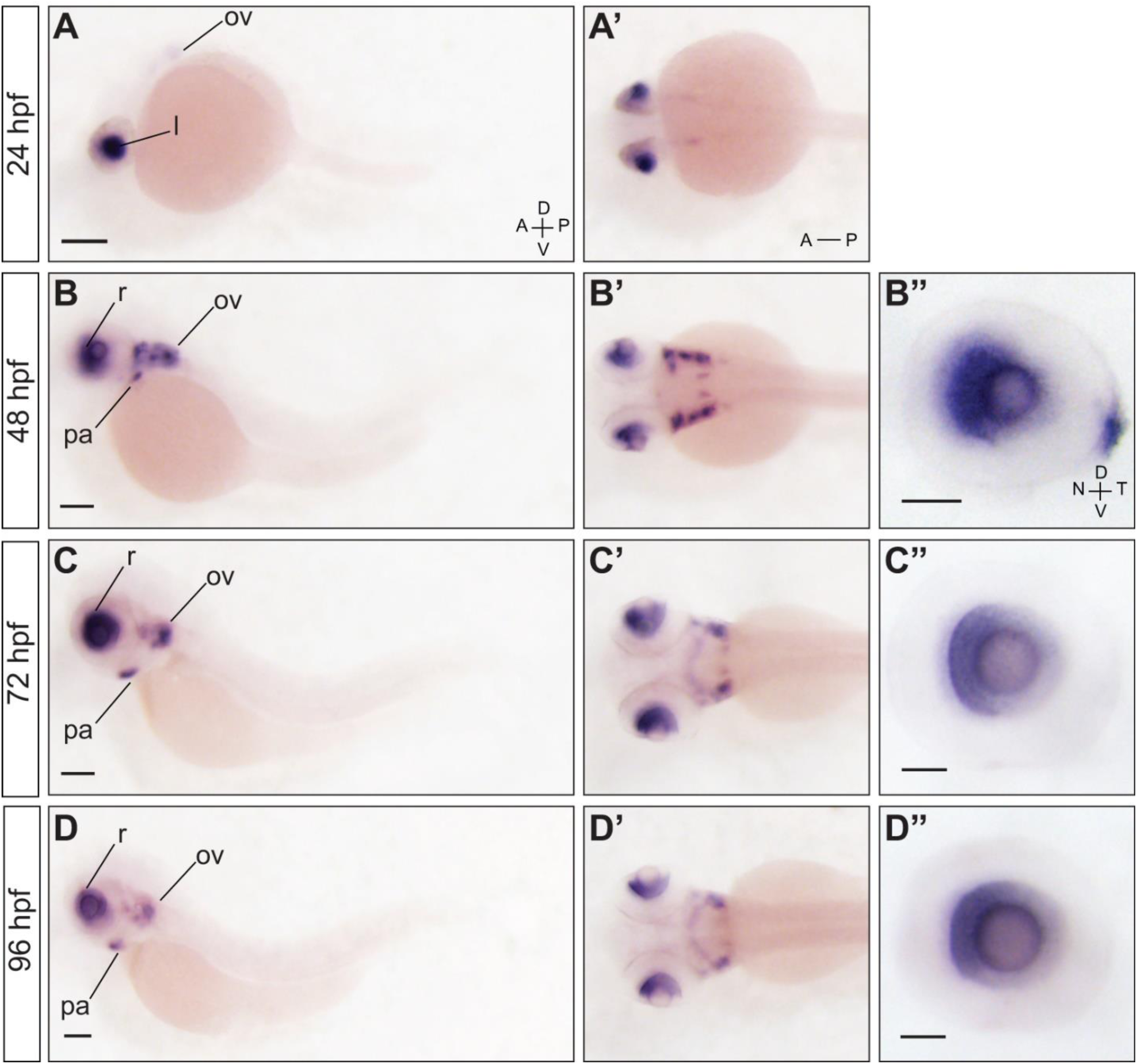
***Hmx4* is expressed in a nasal-high to temporal-low gradient in the retina**. Lateral (**A-D**) and dorsal (**A’-D’**) views of whole embryos stained for *hmx4* by ISH. (**A, A’**) At 24 hpf, *hmx4* is expressed in the developing lens (l) and is also weakly expressed in the retina (r) and otic vesicle (ov). (**B, B’**) At 48 hpf, *hmx4* expression is strongly detected in the anterior retina, the otic vesicle and pharyngeal arches (pa). (**C, C’**) *Hmx4* expression remains consistent at 72 hpf. (**D, D’**) By 96 hpf, *hmx4* remains strongly expressed in the anterior retina and is still detected in the otic vesicle and pharyngeal arches, although to a lesser extent. (**B”-D”**) Lateral views of eyes dissected from embryos stained for *hmx4* by ISH. *Hmx4* has a high-nasal to low-temporal graded expression in the RGC layer of the retina at 48, 72 and 96 hpf. Scale bar: 200 µm in A-D’; 50 µm in E-F.

**Figure 4 - supplement movie 1.** 3D rendered movie of an [*hmx1-En2:cre; RGC:colorswitch*] double transgenic embryo immunostained for TagRFP and EGFP at 4 dpf. Confocal stacks taken from a dorsal view are rotated along the x-axis.To-Pro-3 was used as a nuclear counterstain to visualize the tectal neuropil.

**Figure 5 - supplement figure 1.**
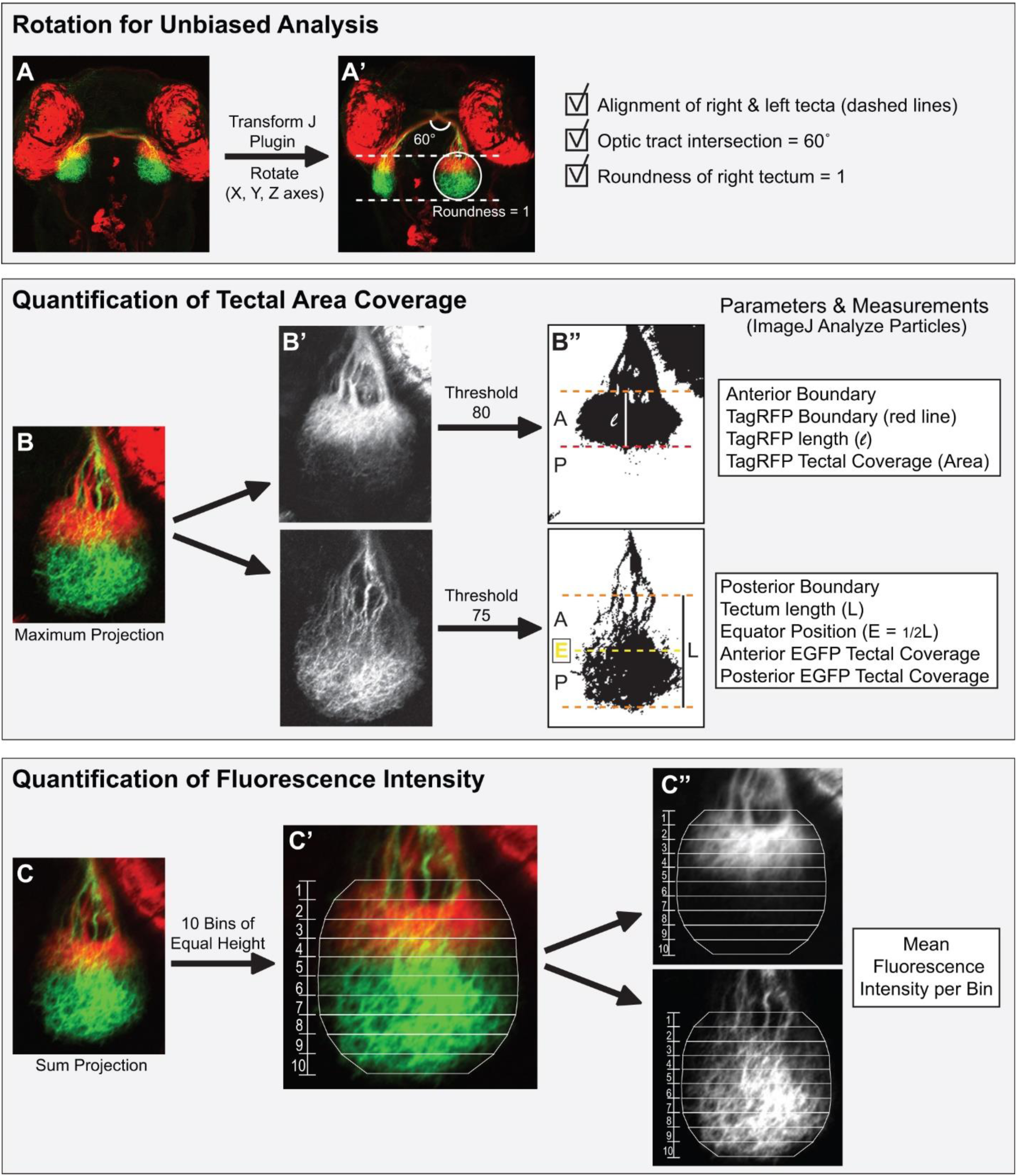
**Visualization and quantification of antero-posterior retinotopic mapping**. (**A, A’**) Confocal stacks taken from a dorsal view were rotated along the X, Y, and Z-axes using the TransformJ plugin in ImageJ to obtain a consistent orientation between embryos. All stacks were rotated so that the left and right tecta were aligned horizontally, both optic tracts intersect at an angle of 60°, and the tectal roundness was equal to 1. (**B-B”**) Quantification of tectal area coverage. (**B, B’**) Rotated stacks were first maximum-projected. (**B”**) TagRFP and EGFP maximal projections were binarized with a threshold of 80 and 75, respectively. The binarized TagRFP maximal projection was used to delineate the anterior tectal boundary, the TagRFP boundary position, and the distance between the anterior tectal and TagRFP boundaries (l). The tectal coverage area of TagRFP-positive temporal axons was measured using the Analyze Particles tool in ImageJ. The binarized EGFP maximal projection was used to delineate the posterior tectal boundary. The total tectum length (L) between the anterior and posterior tectal boundaries was measured, and the Equator (E) was set at ½ the total tectum length. The Analyze Particles tool was used to measure the tectal coverage area of the EGFP-positive nasal axons in the anterior (rostral to E) and posterior (caudal to E) halves of the tectum. (**C-C”**) Analysis of fluorescence intensity at the tectum. (**C**) Confocal stacks were sum-projected. (**C’**) The tectum was divided into 10 bins of equal height using the polygon selection tool, with Bin1 being the anterior-most bin and Bin10 being the posterior-most bin. (**C”**) The mean fluorescence intensity of TagRFP and EGFP signals was measured within each bin using ImageJ.

**Figure 6 - supplement figure 1.**
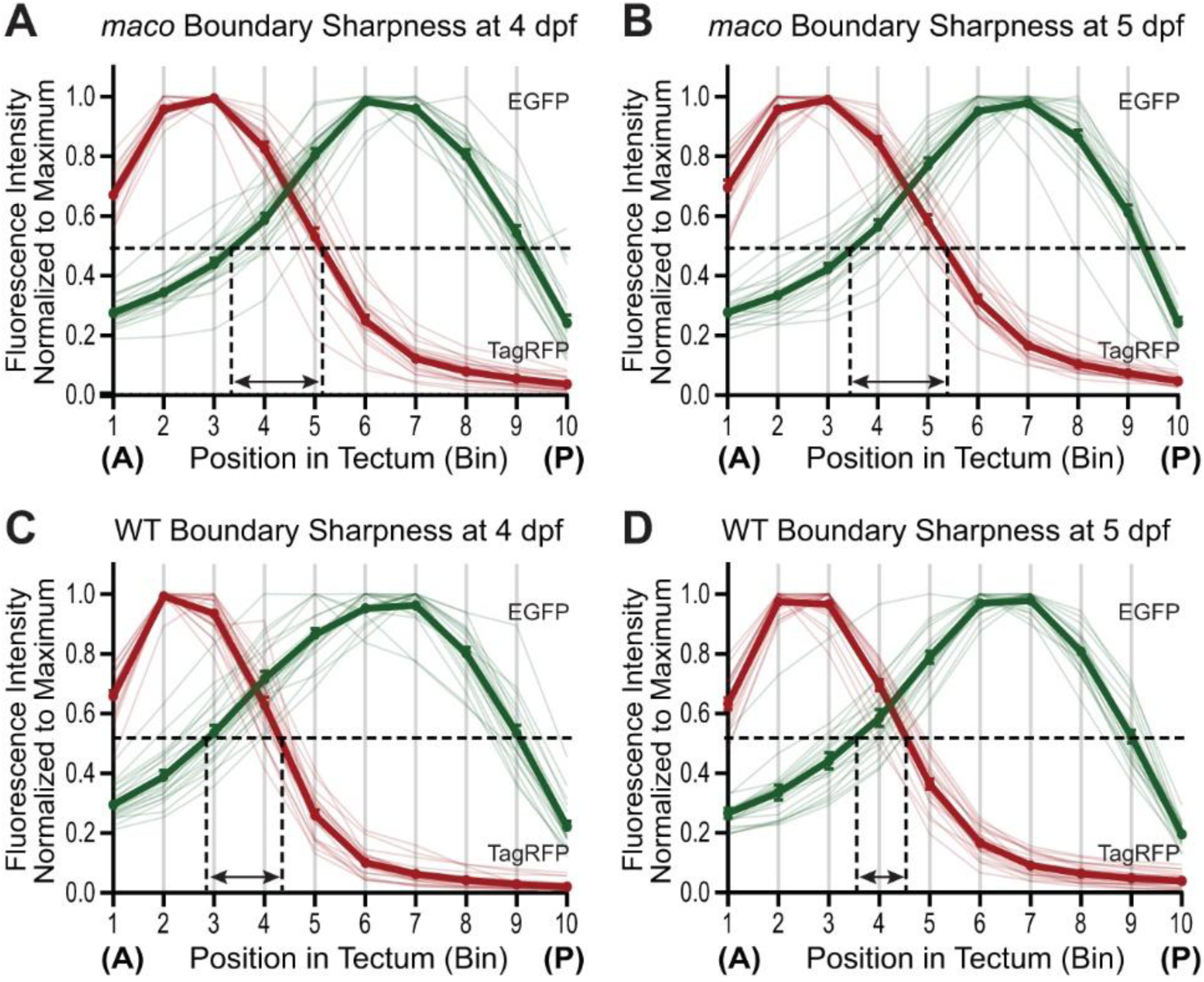
**The antero-posterior retinotopic map does not sharpen over time in *maco* mutants**. (**A-B**) Mean fluorescence intensities of EGFP and TagRFP were normalized to their respective maximum value and plotted along the antero-posterior axis of the tectum. The distance between EGFP_50%_ and TagRFP_50%_ does not change between 4 and 5 dpf in *maco*. (**C-D**) The distance between EGFP_50%_ and TagRFP_50%_ decreases between 4 and 5 dpf in WT siblings, demonstrating that the boundary between EGFP and TagRFP projection domains becomes more precise over time. Data represent mean ± SEM. n = 20 embryos per genotype.

**Figure 6 – supplement figure 2.**
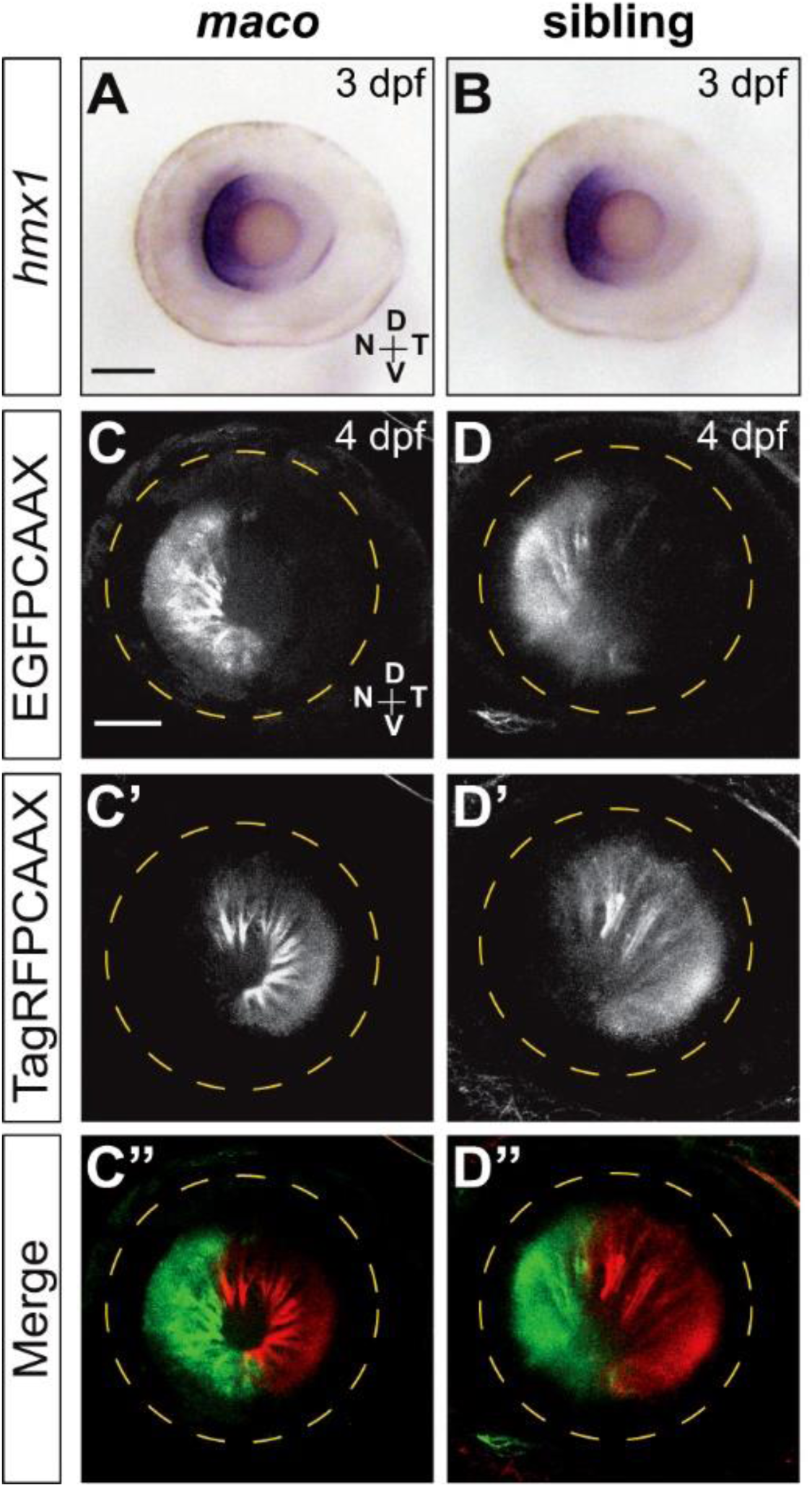
**Retinal patterning is not disrupted in *maco* mutants.** (**A, B**) Lateral views of eyes dissected from *maco* and sibling embryos stained for *hmx1* by ISH at 3 dpf. *Hmx1* expression is restricted to the nasal retina in both *maco* and siblings. (**C-D”**) Eye of a [*hmx1-En2:cre; RGC:colorswitch*] double transgenic *maco* or WT sibling embryo at 4 dpf. (**C, D**) EGFP-positive RGCs are restricted to the nasal retina. (**C’, D’**) TagRFP-positive RGCs are observed in the temporal half of the retina. (**C”, D”**) No differences in EGFP or TagRFP expression in the retina are observed between *maco* and WT. Confocal live microscopy, maximal projections, scale bar: 50 µm.

**Figure 7 - supplement figure 1.**
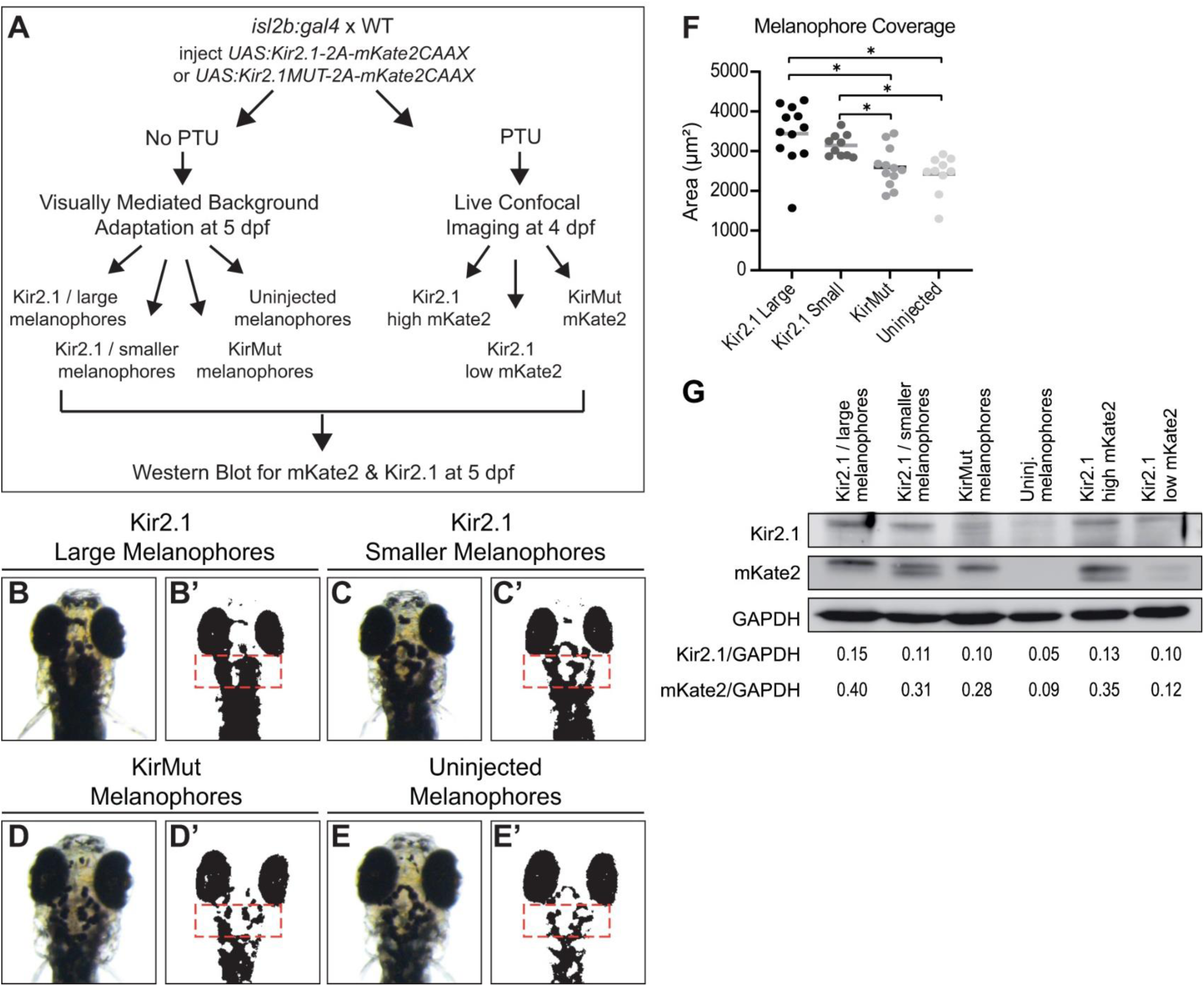
**Expressing Kir2.1 in RGCs blocks larvae’s visually mediated background adaptation**. (**A**) Experimental work-flow to assess the functionality of Kir2.1 transgene in injected embryos. A *UAS:Kir2.1-2A-mKate2CAAX* or a *UAS:Kir2.1MUT-2A-mKate2CAAX* transgene was injected in zygotes from an [*isl2b:gal4*] outcross at one-cell stage. Embryos were divided into two groups: one raised in the absence of PTU for conducting a Visually-mediated Background Adaptation (VBA) assay at 5 dpf, and another raised with PTU for the analysis of mKate2 expression at 4 dpf by confocal microscopy. Based on the VBA assay, embryos were sorted into four groups: embryos expressing Kir2.1 that had expanded pigmentation (“large melanophores”) despite bright illumination, embryos expressing Kir2.1 that had “smaller melanophores”, embryos expressing KirMut, and uninjected embryos as another control group. After confocal imaging at 4 dpf, embryos were sorted into three categories based on the intensity of mKate2 expression: embryos expressing Kir2.1 that had high mKate2 expression (see Material and Methods for high mKate2 expression definition), embryos expressing Kir2.1 that had low mKate2 expression, and embryos expressing KirMut. All embryos were then used for analyzing Kir2.1/Kir2.1MUT and mKate2 expression by Western blot at 5 dpf. (**B-E’**) VBA in embryos expressing Kir2.1 or KirMut in RGCs (thereafter referred to as Kir2.1 or KirMUT embryos) and uninjected embryos. Pictures of embryos in a dorsal view (B, C, D, and E) were binarized using a threshold of 40 (B’, C’, D’ and E’). The area covered by pigmented melanophores was measured in a region caudal to the eyes and rostral to the medulla oblongata (orange box). (**B-C’**) Kir2.1 embryos demonstrated two levels of dark, expanded pigmentation despite bright illumination. (**B, B’**) Some Kir2.1 embryos retained dispersed melanosomes (“large melanophores”) in response to light, indicating a lack of VBA. (**C, C’**) Other Kir2.1 embryos had more restricted but yet abnormally expanded melanin in response to light (“smaller melanophores”), indicating a reduced VBA. (**D-E’**) In contrast to Kir2.1 embryos, KirMut and uninjected embryos showed fully aggregated melanin in response to bright illumination. (**F**) The area covered by pigmented melanophores (pigmentation measured in the region delineated by orange boxes in B’-E’) is significantly larger in Kir2.1 embryos with large or smaller melanophores than in KirMUT or uninjected embryos. Statistical analysis: One-way ANOVA with Tukey’s posthoc test; *p < 0.05, ** p < 0.01, ***p < 0.001. (**G**) Expression of Kir2.1 or KirMUT and mKate2 assessed in the different groups by Western blot at 5 dpf. High expression of Kir2.1 is detected in Kir2.1-expressing embryos with large melanophores and high mKate2 levels. Kir2.1 is also detected at lower levels in Kir2.1-expressing embryos with smaller melanophores or with low mKate2 and in KirMut-expressing embryos. Similarly to Kir2.1, mKate2 is detected in all embryos expressing Kir2.1 or KirMUT but at reduced levels in Kir2.1 embryos with low mKate2, indicating that mKate2 expression can be used as an indicator of Kir2.1 expression. Kir2.1 and mKate2 signal intensities were normalized to that of GAPDH used as a loading control. High levels of Kir2.1 expression correlate with high levels of mKate2 expression. Data represent mean ± SEM. n = 12 Kir2.1 large melanophore embryos, 10 Kir2.1 small melanophore embryos, 12 KirMut embryos, 10 uninjected embryos.

**Figure 7 - supplement figure 2.**
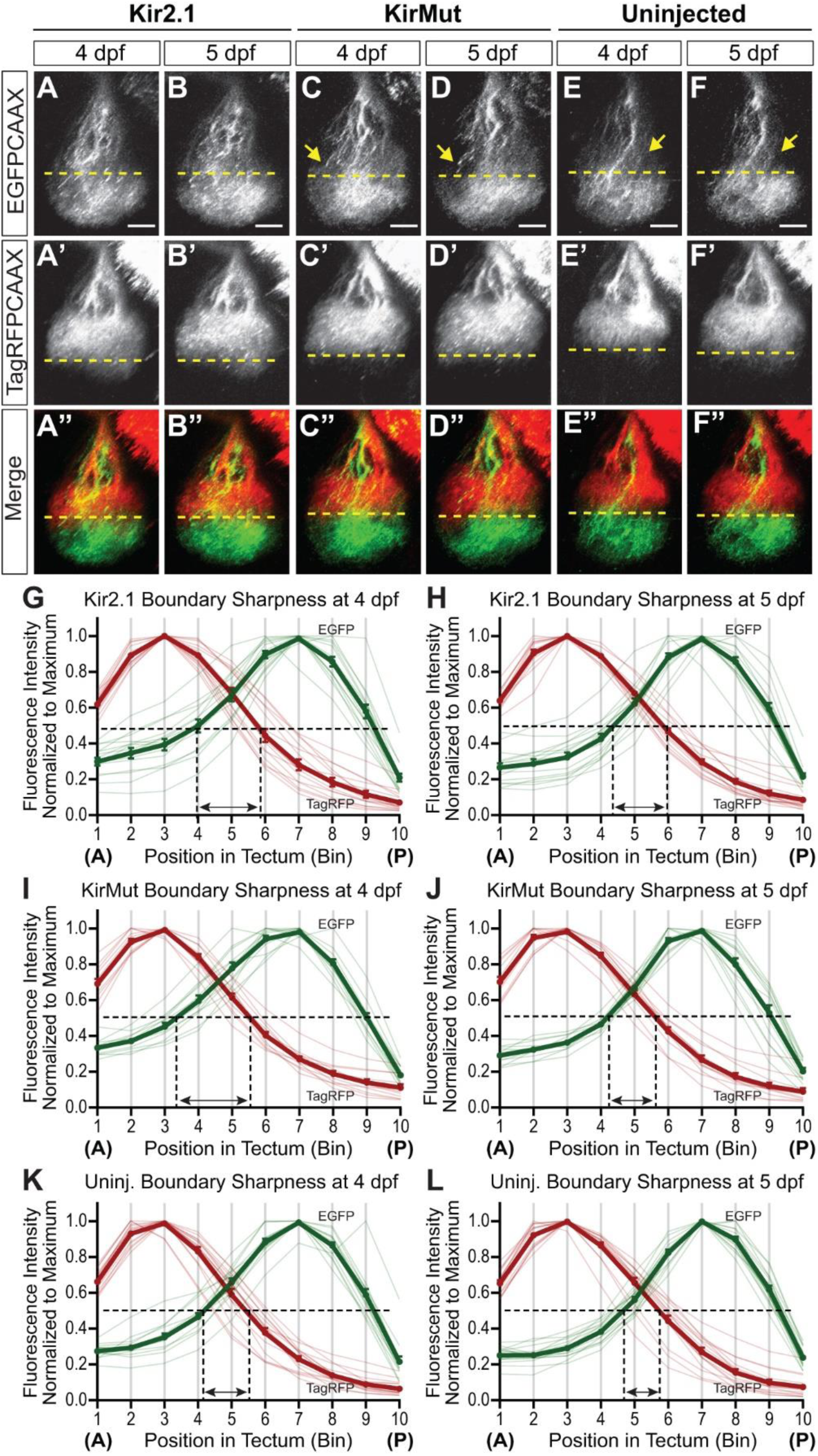
**Blocking neuronal activity in RGCs prevents the sharpening of the antero-posterior retinotopic map**. (**A-F”**) Development of the antero-posterior retinotopic map in Kir2.1 (A-B”), KirMUT (C-D”), and uninjected embryos (E-F”) from 4 to 5 dpf. (**A-F**) EGFP-positive nasal axons innervate the posterior half of the tectum in all three experimental groups. The area covered by nasal axons in the anterior half of the tectum appears to decrease between 4 and 5 dpf in KirMut and uninjected embryos (arrows) but does not seem to change in Kir2.1 embryos. (**A’-F’**) TagRFP-positive temporal axons specifically target the anterior half of the tectum. Confocal microscopy, scale bar: 50 µm. (**G-H**) Mean fluorescence intensities of EGFP and TagRFP were normalized to their respective maximum value and plotted along the antero-posterior axis of the tectum. The boundary sharpness between the temporal and nasal arborization domains (shown as the distance between EGFP_50%_ and TagRFP_50%_) does not change between 4 and 5 dpf in Kir2.1 embryos. (**I-L**) The boundary sharpness between the temporal and nasal arborization domains decreases between 4 and 5 dpf in KirMUT (I, J) and uninjected (K, L) embryos, indicating the formation of a more precise antero-posterior retinotopic map. Data represent mean ± SEM. n = 15 Kir2.1 embryos, 12 KirMut embryos, 15 uninjected embryos.

